# REV-ERBα/β Targeting Transcriptionally Reprograms HIV-Infected CD4^+^ T-Cells for Increased Viral Reactivation but Limited Virion Spread

**DOI:** 10.64898/2026.08.20.746039

**Authors:** Christ Dominique Ngassaki Yoka, Jean-Philippe Goulet, Soumia Khalfi, Debashree Chatterjee, Tomas Raul Wiche Salinas, Yuwei Zhang, Laurence Raymond Marchand, Julie Moreaux, Jonathan Dias, Augustine Fert, Nicolas Chomont, Jonathan Richard, Andrés Finzi, Nicolas Cermakian, Brendan Bell, Jean-Pierre Routy, Laura Solt, Petronela Ancuta

## Abstract

The circadian clock repressors REV-ERBα/β control rhythmic gene expression and inhibit HIV-1 transcription. Whether REV-ERBα/β modulate the HIV-1 replication cycle beyond transcription remains unknown. Here, we demonstrate that memory CD4^+^ T-cells predominantly express the REV-ERBβ isoform *ex vivo* and that T-cell receptor (TCR) triggering downregulates both REV-ERBα/β expression, with levels of REV-ERBα/β mRNA being lower in ART-treated people with HIV (PWH) receiving antiretroviral therapy (ART) compared to people without HIV (PWoH) before/after TCR triggering. In single-round infection, the REV-ERBα/β antagonist SR8278 facilitated HIV-1 reverse transcription, integration, and intracellular HIV-p24 expression, but limited virion release. Moreover, SR8278 downregulated *CCR5* mRNA expression, inhibited R5-tropic HIV-1 replication *in vitro,* and limited viral outgrowth in CD4^+^ T-cells from ART-treated PWH. Finally, genome-wide RNA-sequencing and functional validations revealed HIV-1 restriction/dependency factors that represent novel putative REV-ERBα/β targets. Thus, pharmacological inhibition of REV-ERBα/β uniquely combines a latency reversal activity with the inhibition of progeny virion spread.

Graphical Abstract

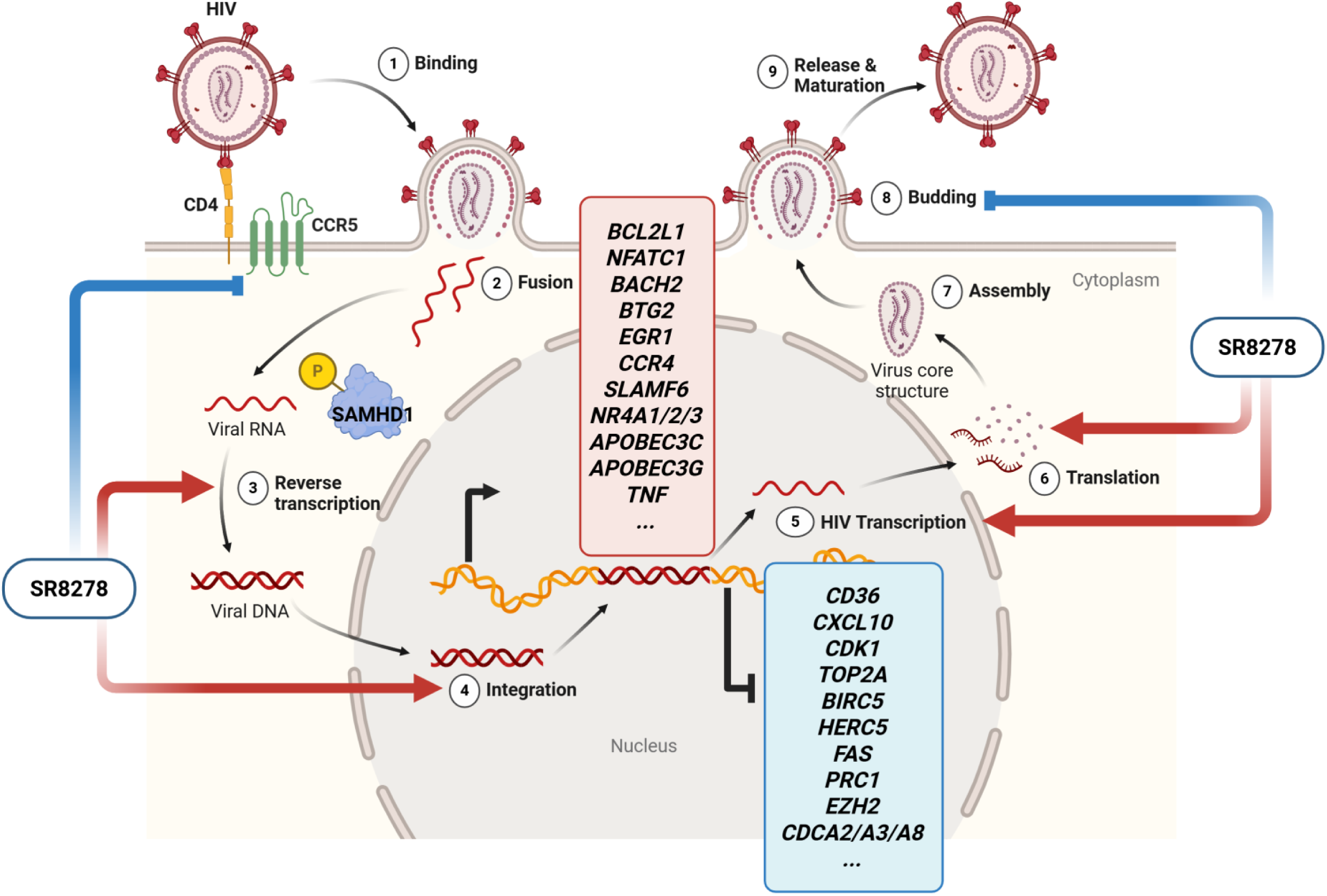

## INTRODUCTION

Antiretroviral therapy (ART) is highly effective at suppressing HIV-1 replication and has transformed a deadly infection into a manageable chronic disease^1–3^. However, it fails to cure HIV-1 from the host, with viral reservoirs persisting despite suppressive ART due to their ability to evade immune-mediated clearance^4,5^. Additional strategies are needed to accelerate reservoir decay and enable sustained viral control in the absence of ART^1,3–9^. One step of the HIV-1 replication cycle that remains untargeted by current ARVs is viral transcription, which depends on the regulatory protein Tat and the host cell transcriptional machinery^10^. While transcriptional latency underlies to the immunological escape of HIV-1 reservoirs, residual HIV-1 transcription during ART contributes to chronic immune activation and increases the risk of co-morbidities^11–14^. This explains current efforts aimed a finding new strategies to modulate HIV-1 transcription by targeting Tat, as well as host cell transcription factors^8,10,15–18^.

CD4^+^ T-cells represent the main targets of HIV-1 infection and constitute the largest reservoir compartment responsible for viral rebound upon ART interruption^1–5,15^. CD4^+^ T-cells are highly heterogeneous in terms of antigenic specificity, functional polarization, pathogenic potential, and tissue-specific homing/residency^15,19,20^. Among these subsets, Th17-polarized CD4^+^ T-cells were identified by our group and others as highly permissive to HIV-1 infection *in vitro*^15^, enriched at mucosal sites of viral transmission^21^, and capable of harboring viral reservoirs in the blood and colon of ART-treated PWH^22–28^. More recently, single-cell RNA sequencing studies revealed persistent HIV-1 reservoirs within CD4^+^ T-cells exhibiting a Th17 transcriptional signature^29,30^. Therefore, understanding molecular mechanisms that regulate HIV-1 permissiveness in Th17 cells may reveal new therapeutic targets for HIV-1 cure interventions.

Our group identified the Th17 master regulator RORC2 (RAR-Related Orphan Receptor C2; RORγt in mice) as a positive regulator of HIV-1 transcription through direct binding to the HIV-1 5’ long terminal repeat (LTR) region^18^. In addition to RORC2, we reported that the transcriptional signature associated with HIV-1 permissiveness in Th17 and Th1Th17 cells includes the circadian clock master regulator component ARNTL (aryl hydrocarbon receptor nuclear translocator-like), also called BMAL1 (Basic helix-loop-helix ARNT-like protein 1)^31,32^. BMAL1 forms a heterodimer with CLOCK (Circadian Locomotor Output Cycles Protein Kaput) that binds E-boxes sequences within the promoters of clock-controlled genes (CCG)^33^. Also, by performing gene set variation analysis (GSVA) of differentially expressed genes (DEG) in Th17 *versus* Th1 cells, we identified the circadian repressor *REV-ERB*α (Reverse strand of *ERBA1*) among top modulated canonical pathways^32^, with REV-ERB exerting a negative feed-back loop on clock genes (CG), such as *RORC2*, *BMAL1*, and *CLOCK*, and other CCG^34^. Studies by Hooper *et al.*, originally demonstrated that Th17 functions are under the control of the circadian clock machinery, with a genetic deficit in REV-ERBα or CLOCK being associated with reduced Th17 frequencies^35^. Further, Solt *et al*. proposed a model in which REV-ERBα inhibits Th17-mediated pathogenicity in an experimental autoimmune encephalitis (EAE) model by preventing the binding of RORγt onto ROR responsive elements (RORE) in the promoter of Th17-specific genes (*i.e., IL17A*)^36^. These findings reveal the rhythmic regulation of Th17 effector functions and support studies on REV-ERBs and other CG/CCG as a potential pharmacological target in autoimmunity^36–39^.

The HIV-1 5’ LTR region contains binding sites for multiple positive and negative regulators of transcription^8,10,16^, including E-box sequences^40,41^, recognized by BMAL1:CLOCK^41,42^ and ROR responsive elements (RORE) that bind RORC2 and REV-ERBs^18,43,44^. Accordingly, previous studies demonstrated direct BMAL1 binding to the HIV-1 promoter and reported circadian variations in cell-associated (CA) unspliced (US) HIV RNA in CD4^+^ T-cells from ART-treated PWH^45,46^. While direct pharmacological targeting of BMAL1 is only beginning to emerge^38^, REV-ERBα/β targeting compounds are already available^47^. In this context, Borrmann et *al*. demonstrated the REV-ERBα/β agonist SR9009 inhibits HIV-1 transcription in TZM-bl cells, primary CD4^+^ T-cells, and macrophages^44^. They also demonstrated that HIV-1 transcription is rhythmic under the control of BMAL1, and RORC2 and REV-ERBα control this rhythmicity by their sequential recruitment onto the HIV-1 promoter^43^. These genes, previously identified as encoding for HIV-1 dependency factors^48,49^, have E-boxes and RORE in their promoters and their expression is modulated by RORC2 inhibitors^43^. Together, this supports the idea that HIV-1 transcription, as well as the transcription of host genes involved in HIV-1 regulation exhibit rhythmic expression patterns and point to REV-ERB as an important negative regulator of these processes.

In this context, we sought to provide new insights into the molecular mechanisms by which the pharmacological targeting of REV-ERBα/β transcriptionally reprograms primary CD4^+^ T-cells for the expression of HIV-1 restriction and/or dependency factors, with an impact on various steps of its replication cycle, beyond viral transcription. Together, our results support the benefits of REV-ERBα/β pharmacological targeting in “*shock and kill*” HIV-1 cure interventions.

## RESULTS

### TCR triggering in CD4^+^ T-cells from PWoH and ART-treated PWH downregulates REV-ERB**α**/**β** mRNA levels, while upregulating Th17 lineage and CCR5 gene expression

The RT-PCR quantification of REV-ERBα/β isoform expression in memory CD4^+^ T-cells from PWoH and ART-treated PWH demonstrated predominant expression of *REV-ERB*β compared to *REV-ERB*α mRNA, both *ex vivo* and following TCR triggering (Figure 1). TCR triggering significantly reduced *REV-ERB*β mRNA expression in PWoH (Figure 1A; Supplemental Figure 1A-B) while increasing *BMAL1, RORC2, IL17A* and *CCR5* mRNA (Supplemental Figure 1C-F). In addition, *REV-ERB*α and *REV-ERB*β mRNA expression was significantly lower in memory CD4^+^ T-cells from ART-treated PWH *versus* PWoH, both *ex vivo* and following TCR triggering (Figure 1B-C). Thus, these results reveal the predominant expression of *REV-ERB*β in human CD4^+^ T-cells, the TCR-mediated downregulation of *REV-ERB*α/β isoforms that coincided with an increased expression of transcripts previously associated with Th17 effector functions and HIV-1 permissiveness^15^.

**Figure 1.**
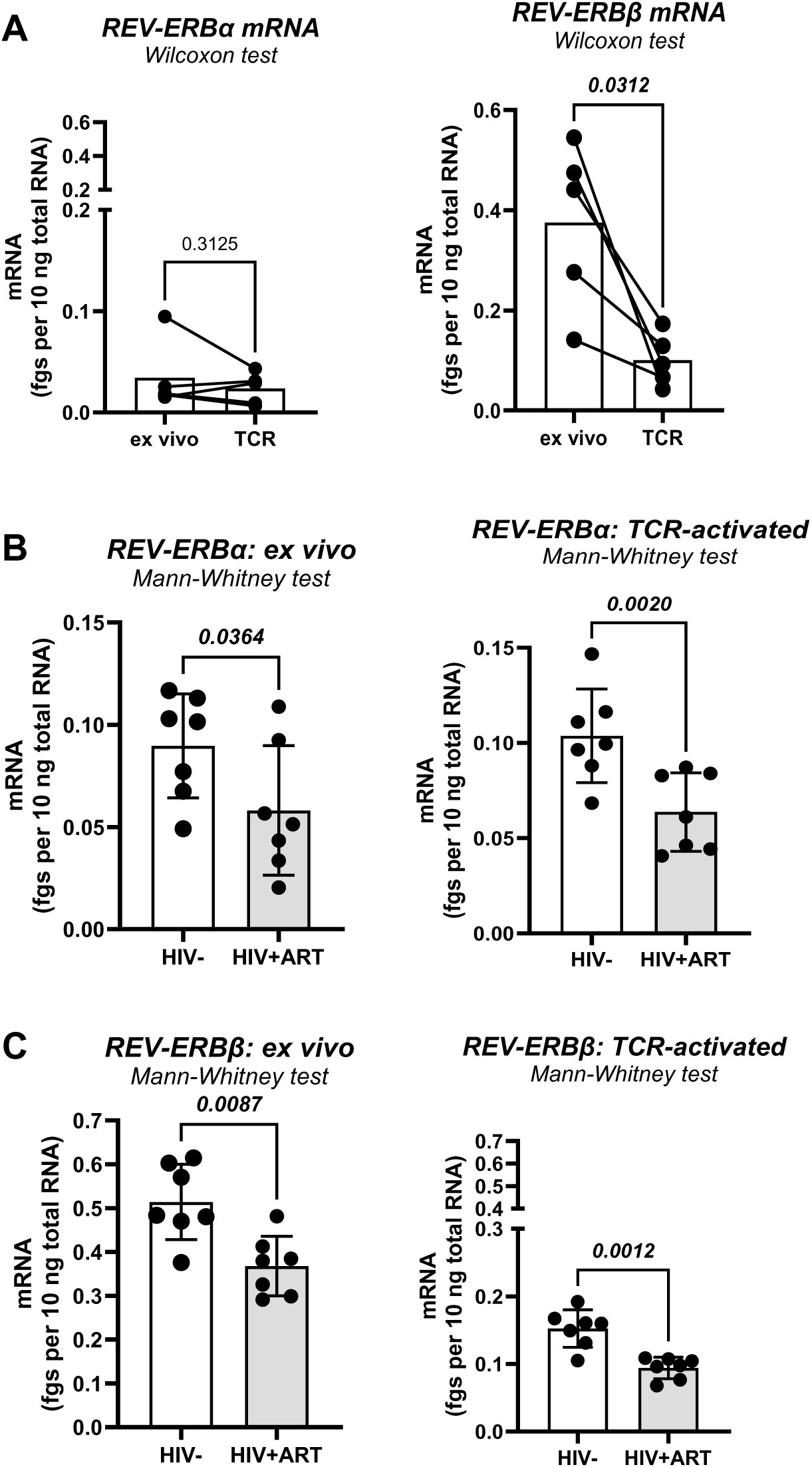
Expression of *REV-ERB*α*/*β mRNA in CD4^+^ T-cells from PWoH and ART-treated PWH. Memory CD4^+^ T-cells from PBMCs of PWoH and ART-treated PWH were isolated by negative selection using magnetic beads (Supplemental Tables 1-2) and stimulated *via* the TCR with CD3/CD28 Abs for 3 days. The *REV-ERB*α and *REV-ERB*β mRNA levels were quantified by real-time RT-PCR in triplicate, relative to a standard curve, and normalized to 28S rRNA levels as a housekeeping gene, allowing to express results in fgs REV-ERB/10 µg 28S rRNA cDNA. **(A**) Shows the levels of *REV-ERB*α and *REV-ERB*β mRNA expression in non-activated (*ex vivo*) *versus* TCR-activated memory CD4^+^ T-cells from PWoH. **(B-C)** Show the levels of *REV-ERB*α **(B)** and *REV-ERB*β **(C)** mRNA expression in non-activated (*ex vivo*, left panel) and TCR-activated memory CD4^+^ T-cells (right panel) from PWoH *versus* ART-treated PWH. Wilcoxon p-values **(A)** and Man-Whitney p-values **(B-C)** are indicated on the graphs.

### Pharmacological REV-ERB targeting modulates *IL17A* and *CCR5* gene transcription

We took advantage of commercially available well-described REV-ERBα/β agonist SR9011^36,37^ and antagonist SR8278^47^ to test their effects on the expression of known CG/CCG in primary memory CD4^+^ T-cells (Figure 2, Supplemental Figure 2). Results show a dose-dependent increase of *REV-ERB*β but not of *REV-ERB*α mRNA expression in the presence of SR8278 (Figure 2A-B), without statistically significant changes observed for *BMAL1* and *RORC2* mRNA (Figure 2C-D). SR8278 significantly decreased *IL17A* at mRNA and protein levels without an effect on *RORC2* (Figure 2D-E and H). Notably, SR8278 decreased the expression of *CCR5* mRNA, without an effect on *CXCR4* mRNA (Figure 2F-G). SR9011 acted similarly to SR8278 in its capacity to increase *REV-ERB*β and decrease *IL17A* and *CCR5 mRNA* expression, with no effect on *REV-ERB*α and *BMAL1* mRNA; however, SR9001 but not SR8278 decreased *RORC2* mRNA expression (Supplemental Figure 2). These results reveal the ability of SR8278 and SR9011 to upregulate the REV-ERBβ expression in CD4^+^ T-cells, while downregulating the expression of Th17 lineage transcripts (*i.e*., IL17A and *RORC2* mRNA) and *CCR5* mRNA expression.

**Figure 2.**
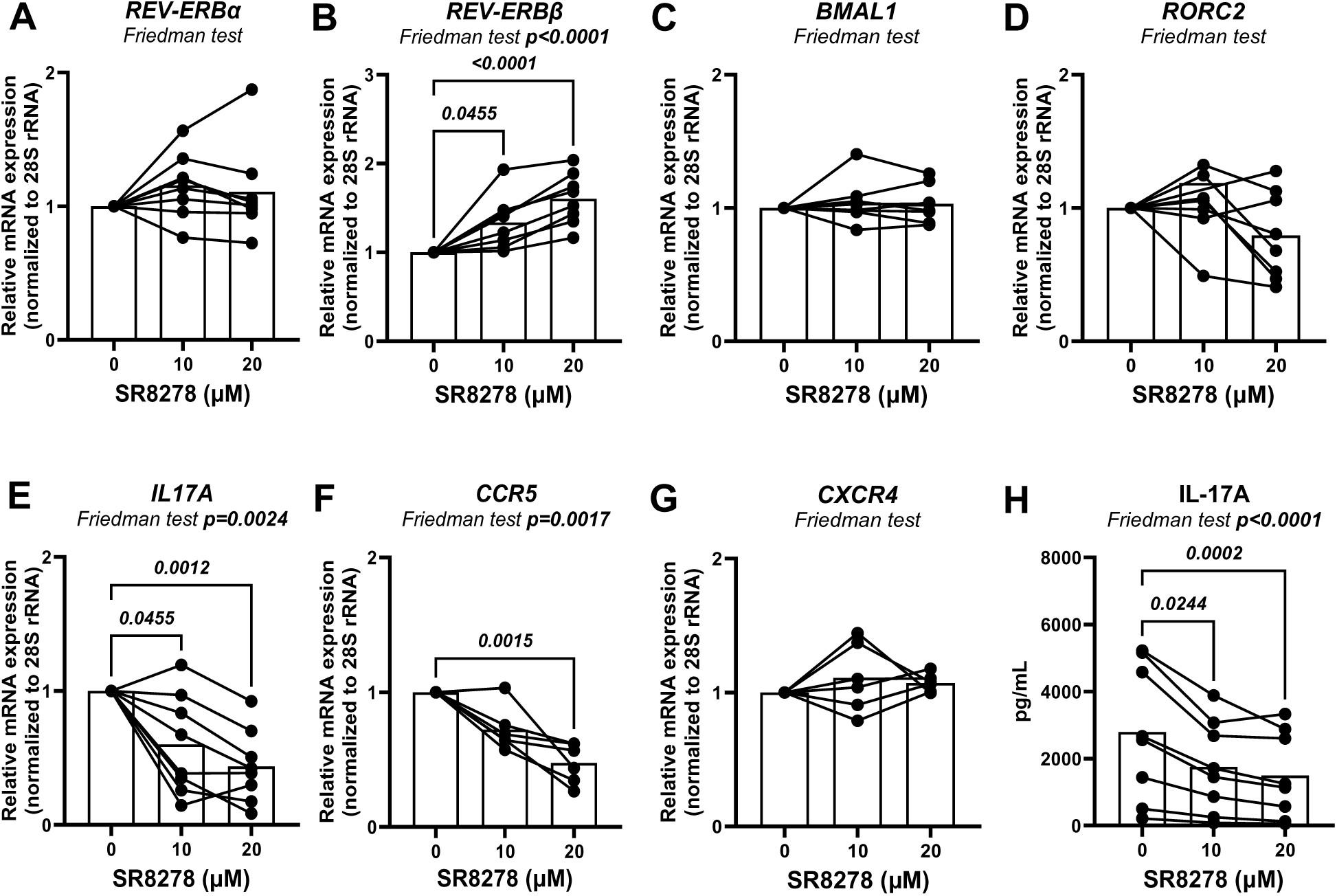
Effects of the REV-ERBα/β antagonist SR8278 on clock genes and HIV-1 co-receptors. Memory CD4^+^ T-cells were isolated from PBMCs of PWoH and gene expression was measured, as described in Figure 1, upon TCR triggering in the presence or the absence of SR8278 (10 and 20 µM), with DMSO used as control. **(A to G)** The levels of *REV-ERB*α and *REV-ERB*β mRNA **(A-B)**, their target genes such as *BMAL1*, *RORC2* and *IL17A (**C-E**)*, as well as *CCR5* and *CXCR4* mRNA (**F-G**). RT-PCR quantification was performed at peak mRNA expression (see results in Supplemental Figure 1), as follows: day 2 post-TCR triggering for *RORC2* and *IL17A*, and day 3 post-TCR triggering for *REV-ERB*α, *REV-ERB*β, *BMAL1, CCR5,* and *CXCR4*. Gene expression was measured in triplicate by one-step real-time RT-PCR relative to 28S rRNA, as a reference gene. Shown are the values of relative gene expression relative to the DMSO condition. IL-17A protein levels were measured by ELISA in cell-culture supernatants harvested at day 3 post-TCR triggering **(H)**. Friedman p-values and uncorrected Dunn’s post-test p-values are indicated on the graphs. Statistically significant p-values (<0.05) are indicated in bold.

### Pharmacological targeting of REV-ERB inhibits R5 HIV-1 replication in CD4^+^ T-cells

To explore the functional consequences of CCR5 mRNA downregulation (Figure 2F), we further investigated the effect of REV-ERB pharmacological targeting on viral replication in memory CD4^+^ T-cells isolated from PWoH using the replication-competent R5 HIV_NL4.3BaL_ virus (Figure 3A). The exposure of memory CD4^+^ T-cells to various doses of SR8278 did not affect cell viability and only slightly reduced the expression of Ki-67, a surrogate marker of cell proliferation^55^, at the highest dose of 10 µM (Figure 3B-C). A statistically significant decrease in HIV_NL4.3BaL_ replication was observed with SR8278 at 10 µM (p=0.001) and at a lower extent at 5 µM (p=0.0552; n=4) (Figure 3D-E). The antiviral effects of SR8278 at 10 µM were confirmed on a larger number of participants (n=10) and coincided with a significant decrease in cell viability measured at day 9 post-infection (Figure 3F-G). Similar to SR8278, exposure to SR9011 significantly reduced Ki-67 expression at 5 µM but not at lower doses, without an effect of cell viability at day 3 post-TCR triggering (Supplemental Figure 3A-B). Also, SR9011 demonstrated antiviral effects at 2.5 and 5 µM (Supplemental Figure 3C-E), but without affecting cell viability at day 9 post-infection (Supplemental Figure 3F).

**Figure 3.**
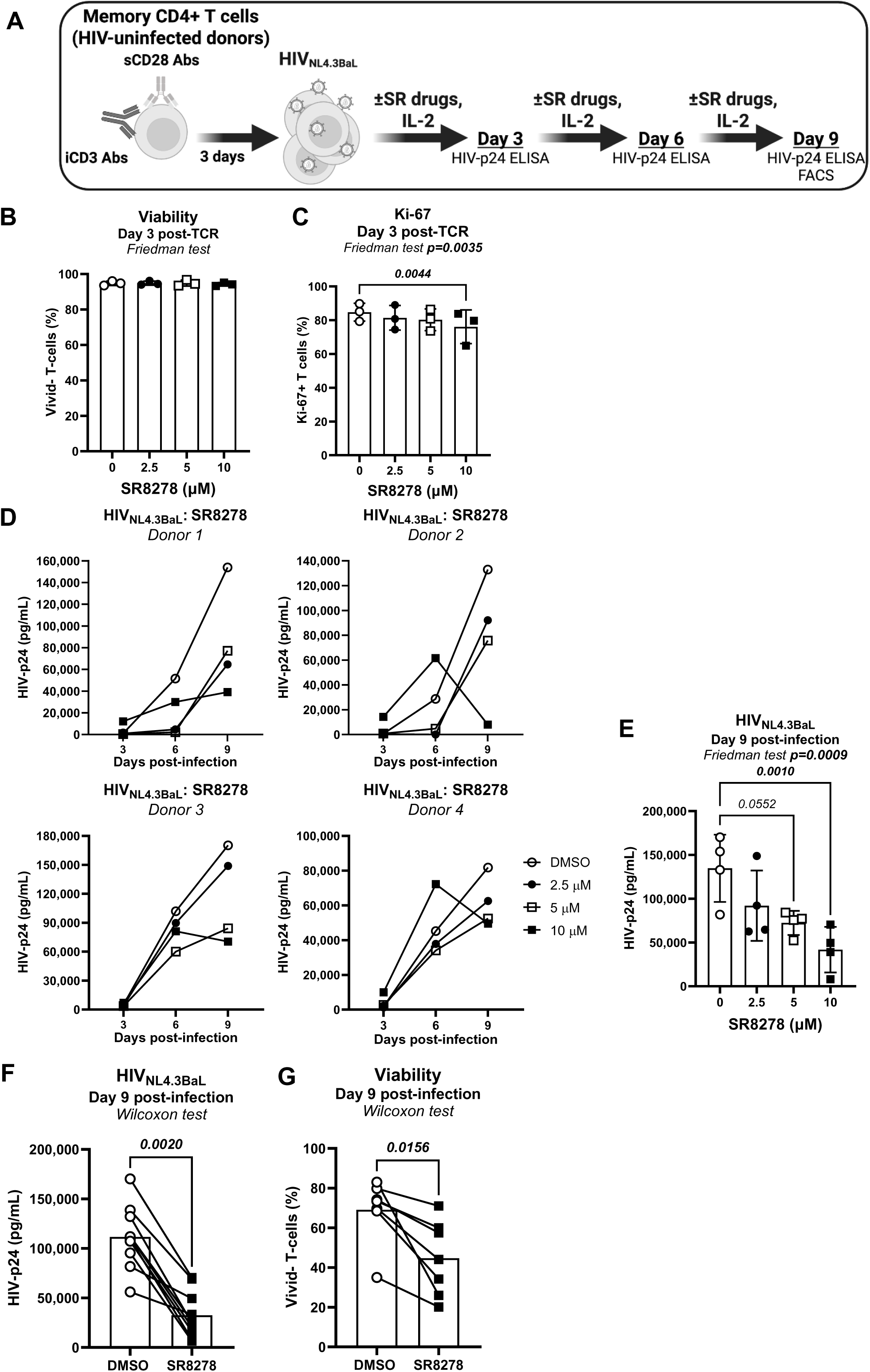
Effects of SR8278 on cell viability and proliferation and HIV-1 replication *in vitro*. **(A)** Shows the experimental flow chart. Briefly, memory CD4^+^ T-cells isolated from PBMCs of PWoH were activated with CD3/CD28 Abs for 3 days, exposed to HIV_NL4.3BaL_ (25 ng/10^6^ cells) and cultured with rhIL-2 (5 ng/ml) in the presence/absence of SR8278 (2.5, 5, 10 µM), with DMSO as control, for up to 9 days. **(B-C)** Show cell survival measured with the viability dye Vivid **(B)** and intranuclear expression of Ki-67, a surrogate marker of cell proliferation **(C)**, at day 3 post-TCR triggering. **(D-E)** Show the kinetics of HIV_NL4.3BaL_ replication in CD4^+^ T-cells in the presence/absence of different concentrations of SR8278 in four participants measured by HIV-p24 ELISA in cell culture supernatants **(D)**, and statistical analysis of HIV-p24 levels at day 9 post-infection(n=4) **(E)**. Friedman p-values and uncorrected Dunn’s post-test p-values are indicated on the graphs. **(F-G)** Show the statistical analyses for HIV_NL4.3BaL_ replication measured by ELISA **(F)** and cell viability measured by FACS **(G)** in CD4^+^ T-cells upon exposure to SR8278 (10 μM) at day 9 post-infection (n=10). Wilcoxon p-values are indicated on the graphs. Statistically significant p-values (<0.05) are indicated in bold.

Thus, SR8278 and SR9011 share a common ability to decrease R5 HIV-1 replication in CD4^+^ T-cells *via* mechanisms involving the interference with cell cycle progression (*i.e.,* Ki-67), consistent with the well-documented role of REV-ERBs as transcriptional repressors^36^ and the reported cytopathic effect of SR8278^50^. In addition, these also results point to CCR5-mediated HIV-1 entry as a mechanism of REV-ERB-mediated viral regulation.

### Pharmacological REV-ERB targeting facilitates post-entry steps of HIV-1 replication cycle

While the antiviral effects of SR8278 and SR9011 could be explained by CCR5 downregulation (Figure 2F, Supplemental Figure 2F), their impact on post-entry steps of HIV-1 replication cycle remain unknown. REV-ERB modulates the transcription of multiple host-cell CCGs^43,47,51^ that may control HIV-1 replication beyond viral transcription. To this end, TCR-activated memory CD4^+^ T-cells were exposed to a single-round VSV-G-pseudotyped HIV (HIV_VSV-G_), which enters cells *via* the LDL receptor^52^, regardless of the HIV-1 receptor CD4 and co-receptors CCR5/CXCR4. HIV-1 replication was assessed three days post-infection by PCR, flow cytometry, and ELISA (Figure 4A). Primers amplifying three different forms of HIV-DNA: *RU5* and *gag* for early and late reverse transcripts, respectively, as well as Alu/HIV-LTR for integrated HIV-DNA were used, as we previously reported^17^. Results in single-round infection demonstrated a significant increased RU5, *gag* and integrated HIV-DNA levels at doses of 10 and 20 µM (Figure 4B), pointing to the facilitation of early/late steps of reverse transcription and integration, despite minor effects on cell viability at 20 but not 10 µM of SR8278 (Figure 4C). Consistently, results obtained by flow cytometry also demonstrated a significant increase in the percentage of GFP^+^HIV-p24^+^ T-cells (Figure 4D-E). In contrast, a significant decrease in virion release was demonstrated by HIV-p24 ELISA quantification in response to SR8278 at 20 but not 10 µM (Figure 4C).

**Figure 4.**
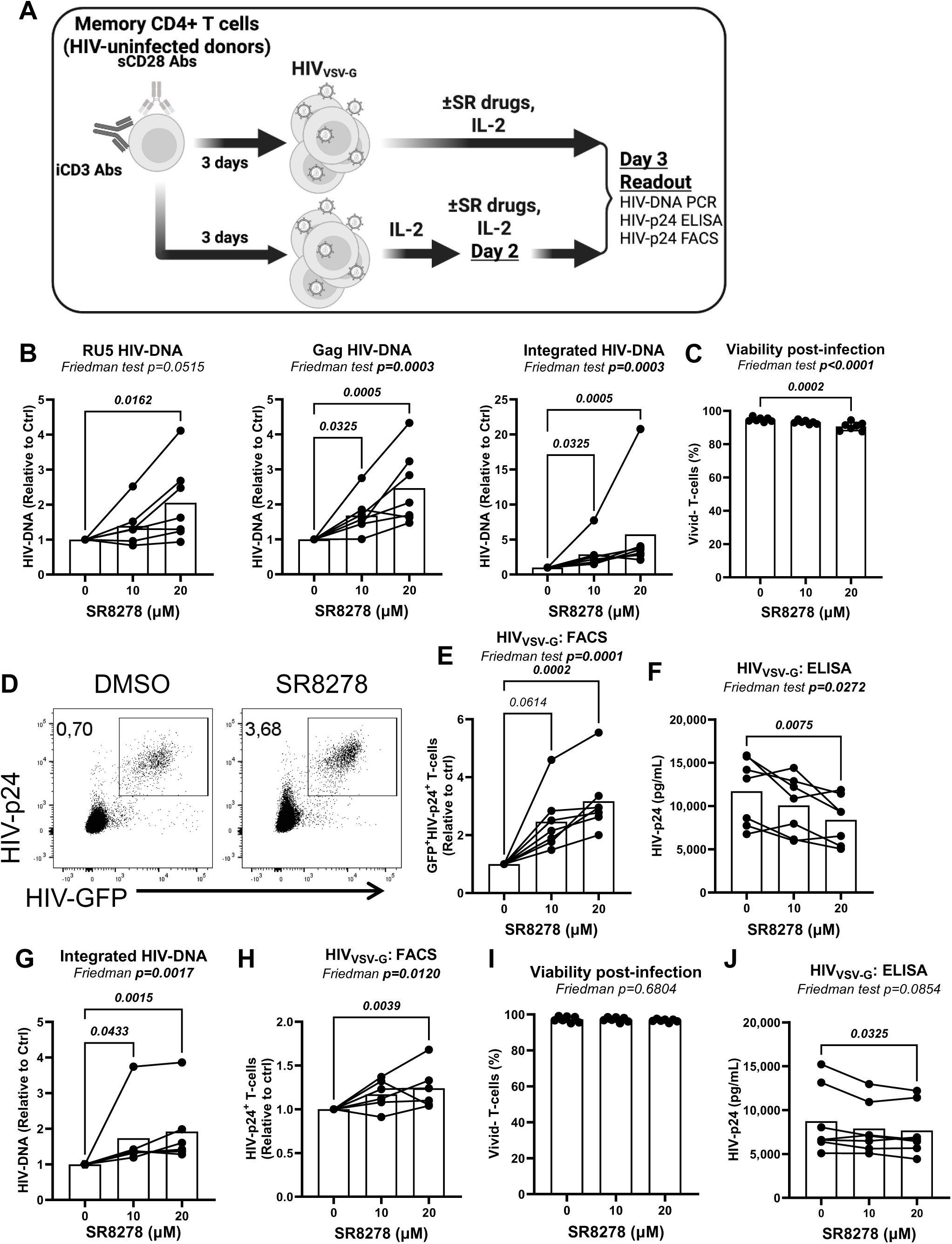
Effects of SR8278 on HIV-1 reverse transcription, integration, translation and virion release. **(A)** Shows the experimental flow chart. Memory CD4^+^ T-cells from PBMCs of PWoH were isolated, stimulated like in Figure 3, exposed to single-round VSV-G-pseudotyped HIV (100 ng/10^6^ cells) and cultured with rhIL-2 (5 ng/ml) for 3 days. In specific wells, SR8278 (10 and 20 µM) was added to cell cultures immediately after infection **(A,** upper side, **B-F)** or at day 2 post-infection **(A,** bottom side, **G-J)**. **(B)** Shows the levels of early (RU5), late (Gag) reverse transcripts and integrated HIV-DNA (copies/10^6^ cells) measured by SYBR Green (RU5) and nested real-time PCR (Gag, Integrated) in CD4^+^ T-cells cultured in the presence/absence of SR8278 relative to DMSO. **(C)** Shows the levels of HIV-p24 in cell culture supernatants measured by ELISA at day 3 post-infection. **(D-F)** Show the levels of intracellular HIV-p24 and GFP expression measured by flow cytometry for one representative donor **(D)**, as well as statistical analysis of intracellular HIV-p24 expression **(E)** and cell viability **(F)** (n=7). **(G-J)** Show the statistical analyses (n=6-7) of the effects of SR8278 added at day 2 post-infection on HIV-DNA integration (copies/10^6^ cells) measured by nested real-time PCR **(G)**, intracellular cellular p24 expression **(H)**, cell viability **(I)** and virion release in supernatants **(J)**. Friedman and uncorrected Dunn’s post-test p-values are indicated on the graphs. Statistically significant p-values (<0.05) are indicated in bold.

To distinguish post-entry from post-integration effects of REV-ERB targeting on HIV-1 replication, in another set of experiments, TCR-activated memory CD4^+^ T-cells were exposed to SR8278 at day 2 post-infection with HIV_VSV-G_ (Figure 4A), when HIV-1 integration is typically maximal, as we previously reported^18^. Under these conditions, unexpectedly SR8278 further increased levels of HIV-DNA integration (Figure 4G), indicative that REV-ERB interferes with post-entry and pre-integration restriction mechanisms, a matter of ongoing research in the HIV field ^48,49^. Subsequently, SR8278 significantly increased the frequency of productively infected CD4^low^HIV-p24^+^ cells at 20 µM, without affecting cell viability (Figure 4H-I). Finally, similar to results in Figure 4C, there was no increase in virion production, but rather a statistically significant decrease in HIV-p24 levels in cell culture supernatants exposed to SR8278 at 20 µM (Figure 4J), pointing to a disconnect between intracellular HIV-p24 expression and virion release.

Similar to SR8278, in this single-round infection model, SR9011 acted post-entry to facilitate HIV reverse transcription and integration (Supplemental Figure 4A), boosted intracellular HIV-p24 expression (Supplemental Figure 4B-C), despite a minor but significant decrease in cell viability (Supplemental Figure 4D), and failed to induce a proportional increase in HIV-p24 levels in cell culture supernatants (Supplemental Figure 4E).

In summary, these results indicate an unexpected disconnection between the proviral and antiviral effects of SR8278 on the post-entry steps of the viral replication cycle, with an increase in HIV-1 reverse transcription, integration and/or translation, without a proportional increase in virion release. These results also reveal unexpected common mechanisms of action for SR8278 and SR9011 on HIV-1 replication *in vitro*.

### Pharmacological REV-ERB targeting promotes SAMHD1 phosphorylation

To investigate mechanisms underlying the enhanced HIV-1 reverse transcription, we examined SAMHD1, a restriction factor that limits HIV-1 reverse transcription in its non-phosphorylated form by limiting intracellular dNTP availability and destabilizing viral RNA, thus preventing the generation of viral cDNA^53^. Western blotting analyses were performed to quantify the expression of total and phosphorylated SAMHD1 in TCR-activated CD4^+^ T-cells exposed to compounds. Both SR8278 and SR9011 increased total and phosphorylated SAMHD1 expression (Supplemental Figure 5), pointing to their potential capacity to limit SAMHD1-mediated restriction on HIV-1 reverse transcription.

### SR9011 but not SR8278 upregulates BST-2 surface expression

To identify mechanisms by which REV-ERBs limit viral release, memory CD4^+^ T-cells infected with HIV_VSV-G_ were analyzed by flow cytometry for the expression of BST2 (Supplemental Figure 6A), an HIV-1 restriction factor that tethers nascent virions at the surface of infected cells^54,55^. BST2 action is counteracted by the viral protein Vpu and productively infected cells typically downregulate BST2 expression relative to uninfected cells^54,55^.^56,57^. CD4 was used as a control (Supplemental Figure 6A), since its expression is also documented to be downregulated on productively infected T-cells by the action of viral proteins Nef and Vpu^58,59^. While CD4 expression on infected and uninfected T-cells was not affected by SR8278 nor SR9011 (Supplemental Figure 6B-C), a statistically significant decrease in BST2 MFI was observed in HIV-p24^+^ T-cells upon exposure to SR8278 at 20 µM (Supplemental Figure 6D). In contrast, the MFI of BST-2 expression increased on both infected and uninfected T-cells in presence of SR9011 at 10 µM, without any effects on CD4 expression (Supplemental Figure 6E). Thus, SR8278 and SR9011 act on BST-2 in different manners, with the increased BST2 expression likely explaining limited viral release in the presence of SR9011 but not SR8278. It is noteworthy that the HIV_VSVG_ construct used in these experiments contains the viral *Vpu* gene that antagonizes BST-2 to facilitate virion release from cell surface^60^. The increase of BST-2 on both uninfected (HIV-p24^-^) and productively infected (HIV-p24^+^) cells suggests that SR9011 likely induces an IFN response, known to enhance BST-2 cell surface levels. Accordingly, IFN treatment has been shown to increase BST-2 surface levels on infected cells despite the presence of Vpu, leading to enhanced virion tethering and consequently higher levels of surface Env^57,60^. In this context, Vpu activity is not directly impaired, as the relative downregulation of BST-2 between infected and uninfected cells is maintained (Supplemental Figure 6A). Rather the overall increase in BST-2 levels likely exceeds the capacity of Vpu, resulting in increased surface BST-2 despite ongoing antagonism^57,60^.

### SR8278 sustains intracellular HIV-p24 expression to the detriment of viral release in TCR-activated memory CD4^+^ T-cells of ART-treated PWH

To evaluate the potential use of REV-ERB compounds as latency reversing agents (LRAs), a viral outgrowth assay (VOA) was performed on memory CD4^+^ T-cells from ART-treated PWH (n=7) cultured in the presence/absence of SR8278 (10 µM) (Figure 5A), using a protocol previously established by our group^17,61,62^. Viral outgrowth was detected in n=4/7 ART-treated PWH tested (Figure 5B). While SR8278 reduced HIV-p24 levels in cell-culture supernatants at day 12 post TCR-triggering for n=3/4 ART-treated PWH (Figure 5B), the intracellular HIV-p24 expression showed no significant reduction induced by SR8278 (Figure 5C). Cell viability remained unaffected throughout the culture period (Figure 5D). Thus, SR8278 sustains the accumulation of HIV-p24 inside the cells to the detriment of virion release, consistent with efficient LRA effects^63^.

**Figure 5.**
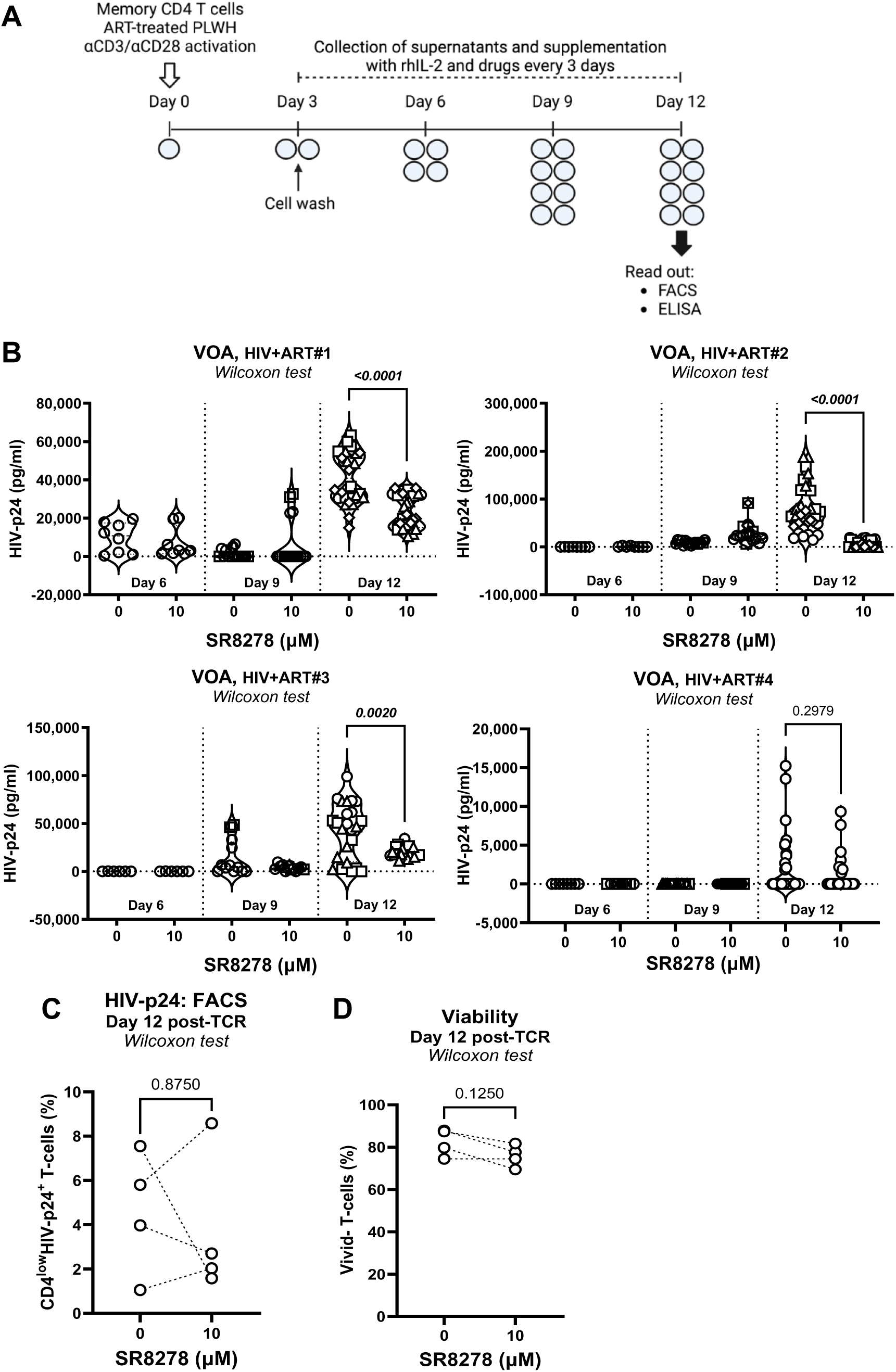
Effects of SR8278 on viral outgrowth in CD4^+^ T-cells of ART-treated PWH. **(A)** Shows the flow chart of the viral outgrowth assay (VOA). Briefly, memory CD4^+^ T-cells isolated from the PBMCs of ART-treated PWH were activated with CD3/CD28 Abs and cultured with rhIL-2 (5 ng/ml) in the presence/absence of SR8278 (10 µM) for 12 days, with media being refreshed and cells split for optimal density (1-2×10^6^ cells/ml) every 3 days. HIV-p24 levels in cell culture supernatants were measured by ELISA. At day 12 post-TCR triggering, cells were harvested and intracellular HIV-p24 expression and cell viability were measured by flow cytometry. **(B)** Shows the levels of HIV-p24 in cell culture supernatants collected at days 6, 9 and 12 post-TCR activation for experiments performed with cells from n=4/7 ART-treated PWH for which viral outgrowth was observed in the DMSO condition. Each symbol indicates a replicate well, with 4, 8, 16, and 32 replicate well generated upon splitting of the 4 original replicates every 3 days. **(C-D)** Show the statistical analyses of intracellular HIV-p24 expression **(C)**, as well as cell viability **(D)**, at day 12 post-TCR triggering. Wilcoxon p-values are indicated on the graphs.

### Transcriptional reprogramming of CD4^+^ T-cells of ART-treated PWH upon REV-ERB pharmacological targeting

To identify additional mechanisms of REV-ERB action, we performed bulk genome-wide RNA sequencing on memory CD4^+^ T-cells from ART-treated PWH (n=7) activated *via* the TCR and cultured in the presence/absence of SR8278 or SR9011 (Figure 6A). Differentially expressed genes (DEGs) were identified based on p-values, adjusted p-values (adj.p), and fold change (FC) ratios (Supplemental Files 1-2) and illustrated in the volcano plot (Figure 6B). Considering significant p-values (<0.05), as well as a FC cut-off of 1.3, we identified 303 (153 upregulated and 150 downregulated) and 642 transcripts (318 upregulated and 324 downregulated) modulated by SR9011 and SR8278, respectively (Supplemental Files 1-2). While SR8278 had a greater impact on gene expression compared to SR9011, a number of 1,085 (p<0.05) and 211 (adj.p<0.05) transcripts were commonly modulated by these two compounds (Supplemental Files 1-2; Supplemental Figure 7A). These results reveal common and distinct mechanisms of action used by SR8278 and SR9011 to transcriptionally reprogram CD4^+^ T-cells in ART-treated PWH, with the most robust effects induced by SR8278.

**Figure 6.**
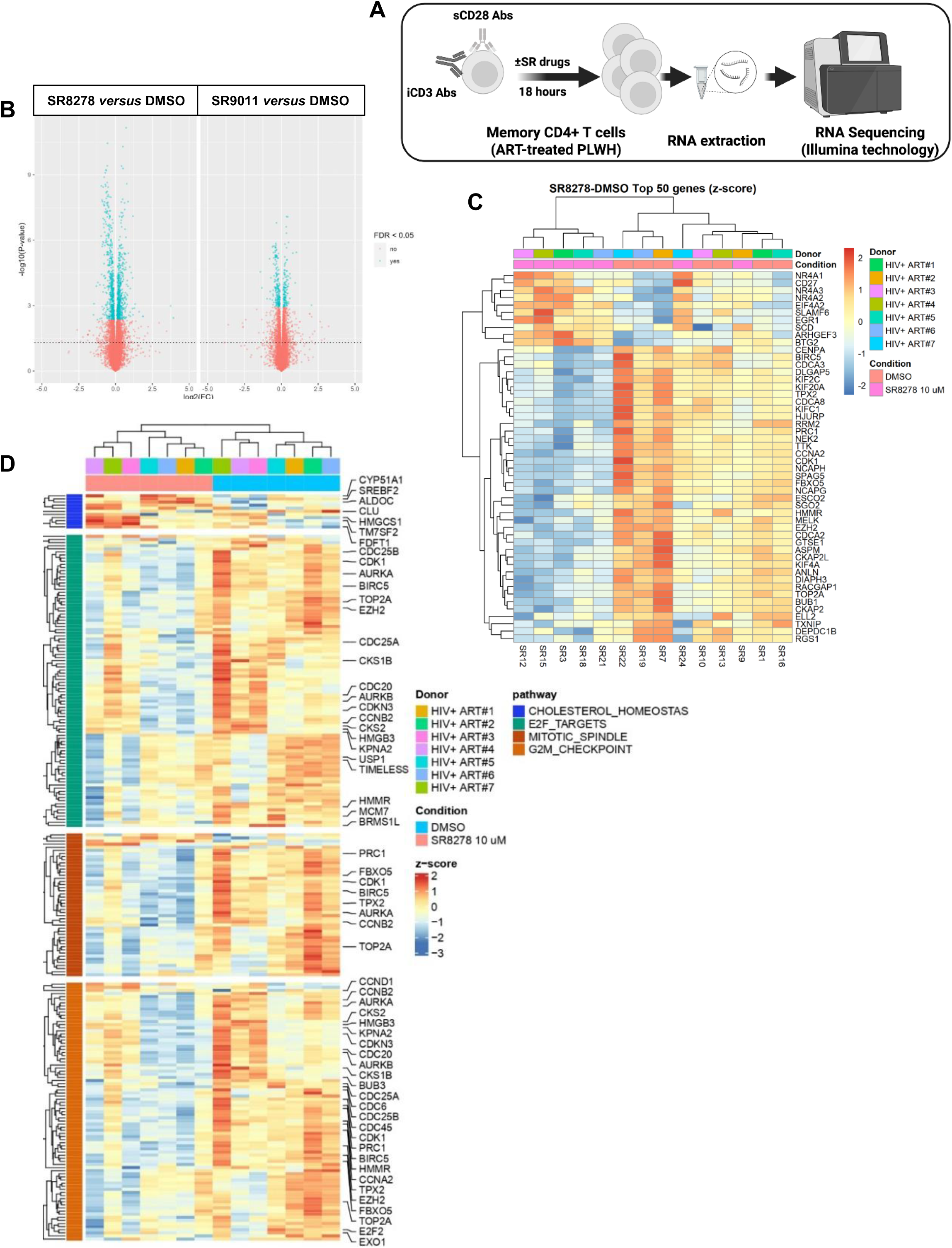
Transcriptional reprogramming of CD4^+^ T-cells upon REV-ERB pharmacological targeting. **(A)** Shows the experimental flow chart of genome-wide bulk RNA sequencing performed using the Illumina technology (25M reads). Memory CD4^+^ T-cells were enriched from PBMCs of ART-treated PWH (n=7) and activated with CD3/CD28 Abs and cultured in the presence/absence of the antagonist SR8278 (10 µM) or the agonist SR9011 (5 µM) for 18 hours. **(B)** Shows the volcano plots indicating differentially expressed genes (DEG) in response to SR8278 or SR9011, with log2 FC (fold-change, 1.3 cut-off) on the x-axis and the negative logarithm of the adjusted p-values for false discovery rate (FDR < 5%) on the y-axis. **(C)** The heatmap depicts the top 50 DEGs based on z-score for the contrast SR8278 *versus* DMSO. **(D)** The heatmap depicts top modulated Hallmark pathways (*i.e.,* cholesterol homeostasis, E2F targets, mitotic spindle, G2M checkpoint) identified by gene set variation analysis (GSVA) with p-values <0.05 for the contrast SR8278 *versus* DMSO.

Among top DEGs, SR8278 upregulated transcripts encoding for nuclear receptors, such as the steroid-thyroid hormone-retinoid receptors (e.g., *NR4A1*, *NR4A2*, *NR4A3*), the anti-proliferative proteins *BTG2*, regulators of the G1/S transition of the cell cycle, as well as factors involved in the regulation of HIV-1 replication (*e.g., EIF4A2*, *SLAMF6*, *EGR1*) (Figure 6C). At the opposite, SR8278 downregulated transcripts encoding for the inhibitor of apoptosis *BIRC5*, cell division cycle-associated genes (*e.g., CDCA2*, *CDCA3*, *CDCA8*), cyclins and cyclin-dependent kinases regulating cell cycle (*e.g., CCNA2*, *CDK1*, *PRC1*), DNA topoisomerase controlling the topologic states of DNA during transcription (*e.g.*, *TOP2A*) and proteins involved in the control of HIV-1 latency (*e.g., PRC1*, *EZH2*) (Figure 6C).

#### Gene set variation analysis (GSVA)

To extract further meaning of the genome-wide transcriptome, GSVA was performed using the UC San Diego/Broad Institute Molecular Signature Database (MSigDB; C2, C3, C5, C7, C8 and hallmark databases; https://www.gsea-msigdb.org/gsea/msigdb/index.jsp). Among the top modulated hallmark pathways identified in GSVA (Supplemental Figure 7B-C), E2F_TARGETS and G2M_CHECKPOINT pathways were downregulated and CHOLESTEROL_HOMEOSTASIS was upregulated by both SR8278 and SR9011 compounds, while MITOTIC_SPINDLE pathway was downregulated by SR8278 but not SR9011 (Supplemental Figure 7D). The preferential effect of SR8278 *versus* SR9011 on mitotic processes may explain its deleterious effects on T-cell proliferation/viability. Transcripts associated with CHOLESTEROL HOMEOSTASIS pathway, involved in multiple metabolic and signaling processes, including HIV-1 replication^64,65^, were mainly upregulated by SR8278 and top DEG included *CYP51A1*, *SREBF2*, *ALDOC*, *CLU*, *HMGCS1*, *TM7SF2*, and *FDFT1* (Figure 6D). The E2F TARGETS include genes that encode for components of the DNA damage checkpoint, repair pathways, chromatin assembly/condensation, chromosome segregation, and the mitotic spindle checkpoint^66,67^. Among transcripts associated with E2F_TARGETS, G2M_CHECKPOINT and MITOTIC_SPINDLE (Figure 6D), we noticed the downregulation by SR8278 of factors involved in DNA replication, cell cycle progression and regulation (*e.g*., *CDC6*, *CDC20*, *CDC25A*, *CDC25B*, *CDC45*), cyclins (*e.g*., *CCNA2*), cycle-regulated kinases (*e.g*., *AURKA*, *AURKB*), DNA topoisomerase (*e.g*., *TOP2A*), polycomb proteins (*e.g*., *EZH2*), proteins regulating cytokinesis (*e.g*., *PRC1*), as well as the downregulation of BIRC5, an inhibitor of apoptosis^68^. Of note, SR8278 downregulated TIMELESS (Figure 6D), a regulator of the circadian clock machinery, as well as cell division and response to cellular stress^69^. Overall, these data indicate that SR8278 disturbs cell cycle progression and regulation, as well as the chromatin landscape that could be in favor of a more efficient HIV-DNA integration, as we have found in (Figure 4B). The regulation of *BIRC5* expression, reported to maintain the survival of HIV-infected CD4^+^ T-cells^68^, suggests an increase of apoptosis in these CD4^+^ T-cells upon SR8278-mediated viral reactivation.

These results reveal the effects of SR8278 on the transcriptional landscape of CD4^+^ T-cells of ART-treated PWH, with consequences on potential specific steps of the HIV-1 replication cycle.

#### Interrogation of the NCBI HIV-1 interactom data base

The NCBI HIV-1 interactom database^70^ was subsequently interrogated for additional data mining using the RNA-Sequencing results in Supplemental Tables 1-2. Among top HIV-1 interactors modulated by SR8278 (*i.e.,* HIV interacts with; HIV upregulates), the upregulated transcripts included those encoding for the pro-inflammatory cytokines *IL2* and *TNF*, the transcription factors *NFATC1*, *NR4A2* (an induced of FOXP3 and Tregs^71^) and *NFKBIA*; the HIV restriction factors *APOBEC3C*, *APOBEC3G*^72^, *TRIM32*, the tetraspanin *CD63*^73^, as well as *BACH2* and *BCL2L1*, involved in HIV integration and T-cell differentiation/survival, respectively^74,75^ (Figure 7A). The top downregulated transcripts included *CXCL10, IL6, CD36* (a mediator of HIV release^76^), *CASP3* (a mediator of apoptosis), *TP73* (a regulator of Tat-mediated HIV transcription), *BIRC5*, *CDK1*, *FAS* (an inducer of apoptosis) and *HERC5* (an inhibitor of HIV mRNA export and assembly^77,78^) (Figure 7A).

**Figure 7.**
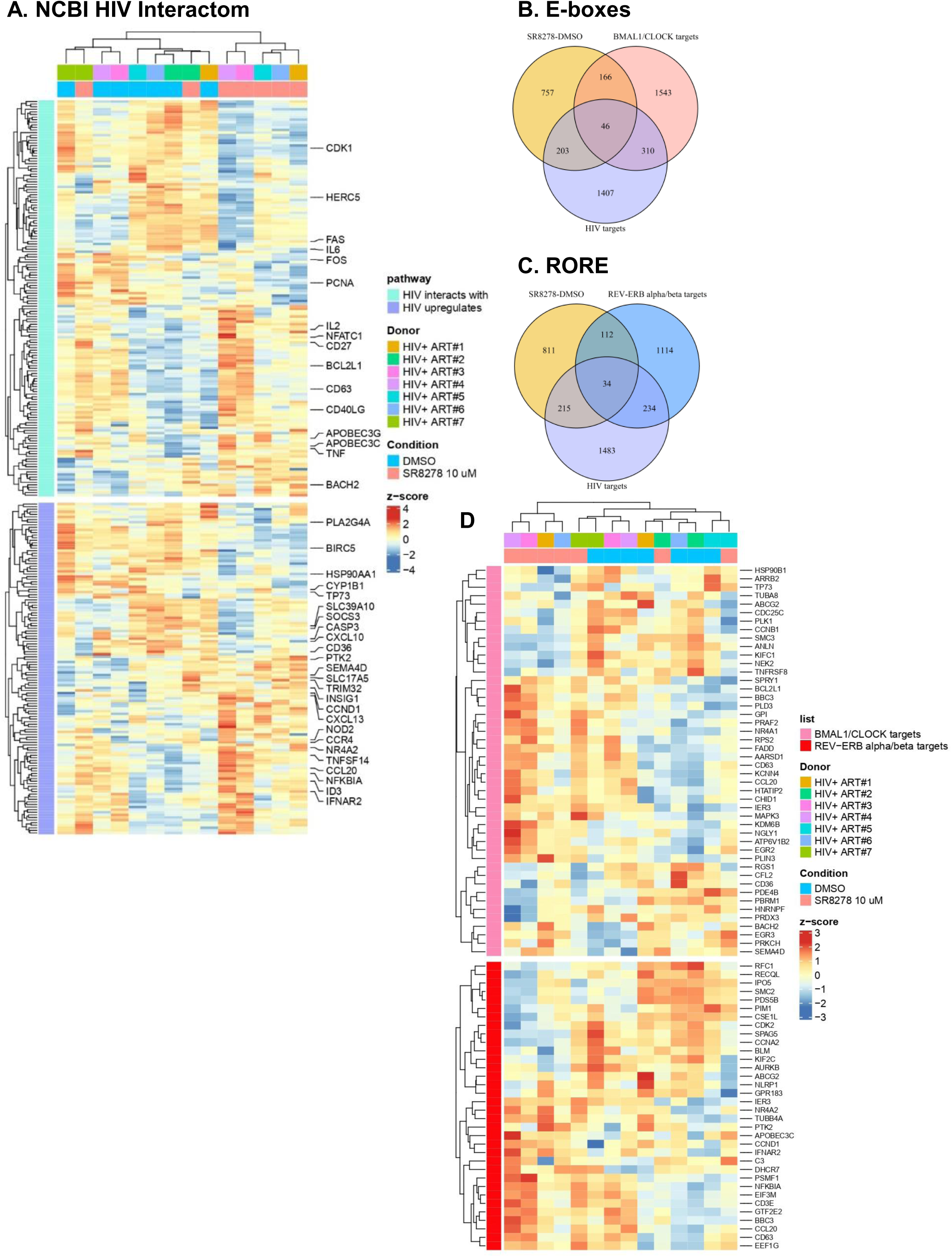
Identification of SR8278-modulated genes included in the NCBI HIV interactom database and expressing E-boxes and ROREs in their promoters. **(A)** RNA-Sequencing experiments were performed as detailed in Figure 6. The heatmap depicts selected transcripts modulated by SR8278 (p-values<0.05; FC cutoff 1.3) matching the lists of genes included on the NCBI HIV interactom database (*i.e.,* upregulates HIV and interacts with HIV terms). **(B-C)** Show the number of genes modulated by SR8278, with/without reported HIV interactions (HIV targets) and with/without E-boxes and ROREs in their promoters. **(D)** The heatmap includes top genes modulated by SR8278, with reported HIV interactions and E-boxes **(top heatmap)** or ROREs **(bottom heatmap)** in their promoters.

### Protein validations of DEGs

The upregulation of *APOBEC3G/C* transcripts (Figure 7A), encoding for a cytidine deaminase reducing the virulence of progeny HIV virions^79^, first caught our attention. Western blotting validations depict a significant increase in APOBEC3G protein expression in CD4^+^ T-cells exposed to SR9011 and SR8278 (Supplemental Figure 8). Moreover, the downregulation of *CD36* transcripts was noted. Its silencing in HIV-1-infected macrophages decreases the amount of newly produced virions released^76^, Flow cytometry validations depict the presence of a small fraction of CD36^+^ CD4^+^ T-cells in the control condition (DMSO) (Supplemental Figure 9A), with the downregulation of CD36 expression induced by SR9011 (%, p=0.0547) and SR8278 (MFI, p=0.0078) (Supplemental Figure 9B-C). Finally, the pro-inflammatory cytokine *TNF* was listed as one of the top upregulated genes in the presence of both SR9011 and SR8278 (adj.p. <0.0001; FC >50) (Supplemental Files 1-2, Figure 7A). Protein validation results confirmed the upregulation of intracellular TNF-α expression by SR8278 (Supplemental Figure 10A-B) and in cell culture supernatants (Supplemental Figure 10C).

### *In silico* identification of direct REV-ERB targets that regulate HIV-1 replication

We sought to identify host factors expressing E-boxes and RORE in their promoters and known as HIV dependency factors among transcripts modulated by SR8278 included in the NCBI HIV interactor database (Figure 7, Supplemental File 2-4). This analysis identified the presence of E-boxes in the promoters of 46 genes modulated by SR8278 and included in the NCBI HIV interactor database, with *BCL2L1*, *MAPK3*, *CCL20*, *CD63*, *HTATIP2*, *NR4A1* and *BACH2* being upregulated, and *CD36*, *CCNB1*, *TNFSF8*, and *HSP90B1* being downregulated (Figure 7B and D; Supplemental File 3). In addition, this analysis revealed the presence of RORE in the promoters of 34 genes modulated by SR8278 and included in the NCBI HIV interactor database, with *NFKBIA*, *CCL20*, *CD63*, *NR4A2*, *PTK2*, *CCND1*, *IFNAR2*, *APOBEC3C* being upregulated, and *CCNA2* and *CDK2* being downregulated (Figure 7C-D; Supplemental File 4).

Together, these results identified novel HIV-1 host factors modulated by SR8278 that represent previously unrecognized BMAL1:CLOCK and RORC2 transcriptional targets modulated by REV-ERB, in line with the expression in their promoters of E-boxes and RORE, respectively.

## DISCUSSION

In this manuscript, we performed immunological, virological and transcriptional investigations on primary CD4^+^ T-cells infected with HIV-1 *in vitro* and viral reservoirs from ART-treated PWH, to gain new insights into the molecular mechanisms by which REV-ERBα/β modulate specific steps of the HIV-1 replication cycle. Using the REV-ERBα/β antagonist SR8278 and agonist SR9011, we identified unexpected discordant effects on distinct stages of the viral replication cycle. Consistent with efficient LRA-like effects^63^, both SR8278 and SR9011 promoted intracellular HIV-p24 accumulation upon infection *in vitro* or in VOA. In addition, both SR8278 and SR9011 facilitated reverse transcription and HIV-DNA integration in a VSV-G-pseudotyped HIV-1 single-cell infection model. These post-entry proviral effects contrasted with a decrease in *CCR5* expression and a reduction in virion release. The genome-wide analysis of transcriptional reprogramming promoted by SR8278 or SR9011 in CD4^+^ T-cells of ART-treated PWH revealed novel mechanisms explaining the intracellular accumulation of HIV-p24 in the absence of a proportional virion release. The subsequent bioinformatics search allowed to list genes encoding for HIV-1 restriction and dependency factors that express E-boxes and ROREs in their promoters, with a potential to be positively regulated by BMAL1:CLOCK and RORC2, respectively, and negatively regulated by REV-ERBα/β. Together, our results support the use of pharmacological REV-ERB targeting in “shock and kill” HIV-1 cure interventions.

The SR8278 and SR9011 compounds used in this study target both REV-ERBα and REV-ERBβ^80^, without allowing discrimination between isoform-specific functions^81^. Nevertheless, an important finding of our study is the predominant expression of REV-ERBβ in human memory CD4^+^ T-cells, suggesting that the effects observed are largely mediated through this isoform. Moreover, another important finding of our study is the downregulation of *REV-ERB*α*/*β mRNA expression following TCR triggering. This raises the hypothesis that TCR triggering might reset or disturb the circadian clock machinery in CD4^+^ T-cells. The TCR-mediated downregulation of transcriptional repressors may explain the increased HIV-1 permissiveness observed upon TCR triggering^82,83^, pointing to a putative role of REV-ERBα/β in restricting HIV-1 infection in quiescent CD4^+^ T-cells. Furthermore, we observed a reduced *REV-ERB*α*/*β mRNA expression in memory CD4^+^ T-cells from ART-treated PWH compared to PWoH. This reduction may explain the state of chronic CD4^+^ T-cell activation largely documented in ART-treated HIV-1 infection^84^. Although the dysregulation of the circadian clock was previously reported during HIV-1 infection^85^, with HIV-1 transcription varying during the day^45,46^. The specific link between REV-ERB and immunological deficits persisting in PWH despite viral suppressive ART remains to be carefully investigated in terms of contribution to HIV-1 reservoir seeding/persistence, as well as disease progression.

SR9011 and SR8278 were previously characterized as agonist and antagonist, respectively^47,80^. Unexpectedly, in our study, both SR9011 and SR8278 shared the ability to increase *REV-ERB*β expression, without an effect on *REV-ERB*α and *BMAL1* mRNA, further supporting the REV-ERBβ-mediated mechanisms of action. Also, both compounds decreased the expression of *IL17A* mRNA and protein, although only SR9011 decreased *RORC2* mRNA expression. Consistently, the RNA-sequencing analysis indicated that SR9011 and SR8278 commonly modulate the expression of certain sets of genes, while showing opposite effects on other sets of genes (Supplemental Figure 8). Thus, our results challenge the classification of SR9011 and SR8278 as exclusive REV-ERB agonist and antagonist, respectively, and rather propose the existence of common mechanisms of action for these two compounds. These results may be explained by multiple structural states taken by these nuclear proteins under the binding of synthetic or natural ligands^86^. Moreover, because of their high dynamism, REV-ERB proteins can respond not only to heme but also to cell redox and gas, which can reverse the repression mediated by the binding of a specific ligand^86^.

Of particular interest, both SR8278 and SR9011 markedly reduced *CCR5* but not *CXCR4* mRNA expression, suggesting that REV-ERB targeting may limit CCR5-mediated HIV-1 entry. This finding is notable given the central role of CCR5 in HIV-1 acquisition/pathogenesis^87^. Indeed, successful HIV-1 cure interventions that target this key HIV-1 co-receptor are reported, including the bone marrow transplantation from donors carrying the protective CCR5Δ32 mutation^88^ and other CCR5 silencing strategies^87^. Although the mechanisms by which REV-ERB regulates CCR5 transcription remain to be determined, our findings identify REV-ERB-targeting compounds as potential modulators of CCR5 expression that warrant further investigation.

SR8278 and SR9011 efficiently modulated HIV-1 replication in CD4^+^ T-cells despite the reduced REV-ERBα/β expression observed following TCR activation. These results may suggest a transient downregulation of REV-ERBα/β protein expression upon TCR triggering, in line with their rhythmic expression^34^. While the SR8278-mediated intracellular HIV-p24 accumulation is consistent with previous studies by Borrmann *et al*. demonstrating an upregulation HIV-1 transcription upon exposure to SR8278^44^, the observed increase in HIV-1 reverse transcription and integration that contrasted with the decrease in viral release was unexpected and previously unrecognized. These discordant pro- and antiviral effects of REV-ERB antagonist are similar to those described for other LRAs (*e.g.,* the PKC inhibitor prostatin)^63,89,90^; and reported by our group for the PPARγ antagonist T0070907^91^ and the AMPK/mTOR modulator metformin^62^. Unlike PPARγ antagonism^91^, REV-ERB antagonism did not boost Th17 effector functions as expected, but rather decreased *IL17A* mRNA expression. The disconnection between the pro- and antiviral actions of these compounds points to a REV-ERB-dependent “*shock and kill*” mechanism of action relevant for HIV-1 cure interventions.

The inhibition of HIV-1 virion release was associated with an increased surface expression of the HIV restriction factor BST2 in the case of SR9011 but not SR8278, indicating discordant mechanisms of action of these two compounds on this step of the viral replication cycle. SR9011 enhances BST-2 levels on both HIV-p24^-^ and HIV-p24^+^ cells, suggesting a viral independent mechanism of BST-2 upregulation, potentially linked to an IFN response. While the viruses used here express Vpu, the increased BST-2 observed on infected cells upon SR9011 treatment likely reflects a condition in which BST-2 induction exceeds the antagonistic capacity of Vpu, resulting in apparent upregulation even in p24+ cells.

The RNA-sequencing analysis oriented us toward cellular factors regulating cell survival and specific steps of the HIV-1 replication cycle, such as integration, transcription and viral release. It is noteworthy that SR8278 upregulated the expression of one transcription factor, *NR4A1*, known to induce apoptosis by its translocation from the nucleus to mitochondria^92^. The upregulation of *NR4A1* mRNA indicates an increased susceptibility to cell death in presence of SR8278, as we observed in our cultures, mainly for high SR8278 doses. Consistently, SR8278 induced the downregulation of *BIRC5*, an inhibitor of apoptosis, essential for the survival of HIV-infected CD4^+^ T-cells^68^. These findings support the idea that REV-ERB antagonism may potentially deplete the pool of productively infected cells *via* NR4A1 and BIRC5-dependent mechanisms under the control of REV-ERB.

The SR8278-mediated increase in *BACH2* expression was also noteworthy given the association between BACH2 and preferential HIV-1 integration sites^93^. Increased *BACH2* expression may therefore contribute to the enhanced HIV-DNA observed following REB-ERB antagonism. However, it is reported that HIV-1 integration near the *BACH2* locus has also been associated with progressive silencing of HIV transcription^94^, consistent with the role of BACH2 as an immune regulator in non-pathogenic Th17 cells^95^. Nevertheless, by modulating *BACH2* expression, REV-ERB may have the potential to regulate HIV reservoir seeding and latency. Whether the SR8278-mediated upregulation of *BACH2* mRNA expression augments HIV-1 transcription by increasing chromatin accessibility for host factors required for HIV-1 gene transcription remains to be investigated. Finally, the GSVA revealed the modulation of pathways (E2F_TARGETS, MITOTIC_SPINDLE, G2M_CHECKPOINT) involved in the progression of the cell cycle, DNA replication and modification of chromatin landscape. Modulation of expression of transcripts such as *PCNA* (proliferating cell nuclear antigen), *CDK1* (cyclin-dependant kinase 1) and *CCND1* (encoding cyclin D1) suggests that SR8278 disturbs essential phases of cell cycle and the chromatin condensation that would be in favor of HIV-1 DNA integration.

The transcriptional changes induced by REV-ERB targeting are consistent with enhanced HIV-1 gene expression. Among the most strongly upregulated genes was TNF, a potent activator of NF-κB signaling and HIV transcription^96^. This suggests that REV-ERB regulates NF-κB, a key factor for HIV-1 transcription and immune activation^10^. Increased expression of NFATC1, another positive regulator of HIV transcription^97,98^, further supports a transcriptionally permissive state. The increased expression of the *NFATC1* gene suggests a positive regulation of NFAT by REV-ERB that increases HIV-1 transcription. SR8278 also upregulated the expression of *NR4A3* mRNA, encoding for a transcription factor associated with stochastic HIV-1 reactivation in latent cell lines and primary CD4^+^ T-cells^99^. This effect of SR8278 on HIV reverse latency is also supported by the downregulation of cellular factors promoting latency, such as PRC1 and EZH2 involved in the epigenetic modifications enforcing HIV-1 silencing^16^. Disruption or downregulation of PRC decreases the methylation of H3K27 and the ubiquitination of H2AK119 in latent cells, leading to HIV-1 silencing^100,101^. Moreover, the increased expression of *EGR1*, which directly drives Tat-dependent HIV transcription^102^, also supports the role of SR8278 in promoting HIV-1 transcription. Finally, *TP73* gene downregulation may increase HIV-1 transcription. In fact, TP73 interacts with Tat and prevents its acetylation, inhibiting the Tat-mediated HIV-1 transcription^103^. The results reported in this article are in line with the literature and push to investigate deeply on the epigenetic modifications caused by REV-ERB antagonism in an indirect manner, at least using latent cell models.

The final steps of the HIV-1 replication cycle consist of virion assembly, maturation and virion release. Our RNA-sequencing analysis identified several host factors implicated in these processes as targets of REV-ERB to interact with HIV-1. This is the case of HERC5 (HECT and RCC1-containing protein 5), which restricts HIV-1 by altering viral mRNA export and the early stage of HIV-1 assembly^78^, supporting the accumulation of intracellular viral components/particles as we observed by flow cytometry. Moreover, the decreased *CD36* mRNA expression, encoding for a class B scavenger receptor which was reported to play a crucial role in HIV-1 release in macrophages^76^, may explain the reduced virion release from productively infected CD4^+^ T-cells exposed to SR8278. Of note, our flow cytometry validations revealed the presence of a small fraction of CD4^+^ T-cells expressing CD36, a fraction that decreased in frequency upon exposure to SR8278 and to a lesser extent SR9011. Furthermore, we found that REV-ERB targeting increased the expression of multiple forms of APOBEC3, including *APOBEC3G* mRNA, encoding for a family of HIV-1 restriction factors counteracted by the accessory protein Vif, and which introduces hypermutations in the genomic HIV-RNA upon incorporation into virions^72^. This may suggest a decrease of HIV-1 infectivity in presence of SR8278. Although the RNA-sequencing results brought interesting insights on genes differentially regulated by SR8278, these results did not allow us to identify a unique mechanism of action, thus supporting the pleiotropic functions of genes regulated by REV-ERB.

Finally, in our quest for genes directly modulated by REV-ERB with pro- and antiviral functions we identified SR8278-modulated genes included in the NCBI HIV gene list for keywords “upregulates” and “interacts” and expressing in their promoters E-boxes (*e.g., BCL2L1*, *MAPK3*, *CCL20*, *CD63*, *HTATIP2*, *NR4A1*, *BACH2*, *CD36*, *CCNB1*, *TNFSF8*, *HSP90B1*) and RORE (*e.g., NFKBIA*, *CCL20*, *CD63*, *NR4A2*, *PTK2*, *CCND1*, *IFNAR2*, *APOBEC3C*, *CCNA2*, *CDK2*) that can serve as putative binding sites for BMAL1:CLOCK and RORC2, respectively (Figure 7).

In summary, our results reveal the pleiotropic effects of REV-ERB pharmacological targeting on multiple steps of the HIV-1 replication cycle. REV-ERB modulation reduced CCR5 expression and virion release while simultaneously enhancing HIV reverse transcription, integration, transcription and intracellular viral protein expression in CD4^+^ T-cells (Graphical Abstract). Given that REV-ERB is a transcriptional repressor, our results support the possibility that REV-ERB compounds interfere with chromatin condensation, thus favoring HIV-DNA integration and then promoting HIV-1 transcription by specific transcription factors. Finally, based on these findings, we identify REV-ERBs as novel targets for potent LRAs and emphasize the need to design clinical-grade agonists/antagonists for «*shock and kill*» HIV-1 cure strategies.

## LIMITATIONS OF THE STUDY

This study has several limitations. First, REV-ERB function was investigated using pharmacological compounds rather than genetic approaches in an effort to characterize new drugs to be used in HIV cure interventions. Second, although we quantified HIV-1 translation, we did not directly measure viral transcription, which was previously shown to be regulated by REV-ERB through direct interactions with the HIV-1 promoter^40,43^. Our readouts may have been partially impacted by the fact that SR8278 decreased cell viability. We argue this effect is related to the ability of REV-ERB to modulate cell death pathways, such as apoptosis in infected cells, as reflected by our RNA-sequencing data. In addition, the VOA was performed on a limited number of ART-treated participants, with detectable outgrowth observed in only a subset of participants. Further studies should be extended to a large number of ART-treated PWH. Moreover, the latency-reversing effects of REV-ERB compounds should be tested using other techniques such as the HIV-Flow^104^ and/or HIV Flow-FISH assay^105^. Furthermore, whether the CD4^low^HIV-p24^+^ T-cells are recognized by HIV-Env broadly neutralizing Abs (bNAbs)^62^ and/or targeted by Abs-dependent cellular cytotoxicity (ADCC) for depletion^106^ remains to be determined *in vitro* and in preclinical animal studies. Finally, validations are needed to determine the direct regulation by REV-ERB of genes encoding for HIV-1 restriction/dependency factors.

## Authors’ contribution

C.D.N.Y. performed the majority of the experiments, analyzed the data, prepared the figures and wrote the manuscript. J.P.G. performed the analysis of RNA sequencing results and prepared figures. S.K., D.C. and T.R.W.S. contributed to experimental design. Y.Z. participated in research design and performed preliminary experiments in support of the study. L.R.M., J.M., J.D. and A.Fe. set up research protocols and performed experiments. N.Ch, J.R., A.F., N.Ce., B.B., and J.P.R. contributed to experimental design, provided protocols, analyzed results and manuscript revisions. N.Ch., J.R., A.Fi., N.Ce., and B.B. provided protocols, contributed to clinical data for results analysis and manuscript revisions. J-P.R. recruited study participants, provided access to leukapheresis and ART-PWH and Pw/oH, and contributed to manuscript revisions. L.S. contributed to the research design, provided reagents for preliminary experiments and analyzed data. P.A. designed the study, provided supervision/training, analyzed results, prepared figures, and wrote the manuscript. All authors revised the manuscript and approved its submission.

## Acknowledgement

The authors thank Dr. Gael Dulude, Dr. Dominique Gauchat and Philippe St Onge (Flow Cytometry Core Facility, CHUM-Research Center, Montréal, QC, Canada) for technical support in flow cytometry; Dr Olfa Debbeche (Biosafety Level 3 Core Facility CHUM-Research Center, Montréal, QC, Canada); Mario Legault (FRQ-S/AIDS and Infectious Diseases Network; Montreal, QC, Canada) for help with ethical approvals and informed consents; Josée Girouad and Angie Massicotte (McGill University Health Centre, Montreal, QC, Canada) for their key contribution to blood collection and clinical information from PWH and negative study participants. The authors also thank Dr. Roger Pomerantz (Thomas Jefferson University, Philadelphia, PA, USA) for providing the NL4.3BaL HIV plasmid, Dr. Michel Tremblay (Université Laval, Quebec, QC, Canada) for providing the HIV-p24 antibodies producing hybridoma and Dr Andrés Finzi for providing anti-envelope antibodies. Finally, the authors acknowledge the key contributions of all study participants for their precious gift of leukapheresis essential for this study.

## STAR METHODS RESOURCE AVAILABILITY

### Lead contact

Further information and requests for resources and reagents should be directed to and will be fulfilled by the lead contact Petronela Ancuta.

## MATERIALS AVAILABILITY

This study did not generate unique reagents.

## Data and code availability

The entire RNA-Sequencing data set and the technical information requested by Minimum Information About a Microarray Experiment (MIAME) are available at the GEO database under accession GSE253420.

## EXPERIMENTAL MODEL AND STUDY PARTICIPANT DETAILS

### Ethics statement

This study was performed on leukapheresis collected from ART-treated PWH and PWoH, while respecting the principles included in the Declaration of Helsinki. The study was approved by the Institutional Review Board of the McGill University Health Centre and the CHUM-Research Centre, Montreal, Quebec, Canada [2023-11299 (22.267)]. All study participants signed informed consents and agreed with the publication of the results.

## METHOD DETAILS

### Study participants

Leukapheresis samples were collected from PWoH and ART-treated PWH, with plasma VL < 40 HIV-RNA copies/mL (Supplemental Tables 1-2). Study participants were recruited at the McGill University Health Centre and Centre Hospitalier de l’Université de Montréal (CHUM, Montréal, Québec, Canada). PBMCs were isolated from leukapheresis by gradient centrifugation using the lymphocyte separation medium (Wisent, Saint-Jean-Baptiste/Canada), and preserved frozen in 10% DMSO (SIGMA, St. Louis/United States) and 90% fetal bovine serum (FBS; Wisent, Saint-Jean-Baptiste/Canada) in liquid nitrogen.

### Cell enrichment and TCR activation

Memory CD4^+^ T cells were enriched from PBMCs by negative selection using magnetic beads (EasySep™ Human Memory CD4^+^ T Cell Enrichment Kit, STEMCELL Technologies, Vancouver, BC, Canada). Cell purity (>95% purity) was assessed upon staining with fluorochrome-conjugated CD3, CD4, CD45RA antibodies (Supplemental Table 3) and flow cytometry analysis (see below). Cells were TCR-activated using immobilized anti-CD3 and soluble anti-CD28 antibodies (1 µg/mL, BD Pharmingen, San Diego, CA, USA) for 3 days and used for subsequent experiments.

### Virus stocks and *in vitro* infection

Two types of viruses were used to perform *in vitro* HIV-1 infection: replication-competent CCR5-tropic NL4.3BaL (HIV_NL4.3BaL_) and single-round vesicular stomatitis virus (VSV)-G-pseudotyped-HIV-1 (HIV_VSVG_). HIV_NL4.3BaL_ is a NL4.3-based provirus expressing the BaL envelope, kindly given by Michel Tremblay (Université de Laval, Québec, Canada) and originating from Dr Roger J Pomerantz (Thomas Jefferson University, Philadelphia, Pennsylvania, USA). The HIV_VSVG_ stock was produced from a plasmid encoding for an env-NL4.3-based provirus, with *gfp* in place of *nef*, and another plasmid encoding for VSV-G envelope. HIV-1 stocks were generated by transfection in 293T using X-tremeGENE HP DNA Transfection Reagent (Roche Diagnostics, Mannheim, Germany) as we previously described. TCR-activated memory CD4^+^ T-cells were exposed to HIV-1 (25 ng HIV-p24/10^6^ cells) for 3 hours at 37 °C. After a washing step to remove the unbound virions, cells were cultured in the presence of IL-2 (5 ng/ml, R&D Systems, Minneapolis, MN, USA), and in the presence/absence of SR9011 or SR8278 (concentrations indicated in Figure legends) up to 9 days. The media was refreshed every 3 days. Viral replication was evaluated by flow cytometry and ELISA (see below). In parallel, cells were harvested at day 3 post-infection for PCR-based HIV-DNA quantification as described below.

### Chemical synthesis and reagents

SR9011 and SR8278 have been previously described^80,107^.

### Flow cytometry analysis

For surface staining, cells were harvested and washed using a FACS buffer (1X PBS, 10% FBS, EDTA 2mM, 0.2% sodium azide) and incubated with fluorochrome-conjugated antibodies (Supplemental Table 3) for 30 minutes at 4 °C. For intracellular staining, HIV-1 infected cells were permeabilized and fixed using the BD Cytofix/Cytoperm fixation/permeabilization Solution kit (according to the manufacturer’s recommendations. For intranuclear staining, CD4^+^ T cells were permeabilized and fixed using the eBioscience™ Foxp3/Transcription Factor Staining Buffer Set (according to the manufacturer’s recommendations. Dead cells were excluded by using the live/dead fixable aqua dead cell stain kit. Samples were acquired by flow cytometry using the LSRIIA and FORTESSA cytometers powered by BD FACS Diva software (BD Bioscience, San Jose, CA, California). Finally, flow cytometry data were analyzed using BD Flowjo (Tree Star, Inc., Ashland, Oregon, USA).

### ELISA

Soluble HIV-p24 levels were quantified in cell culture supernatants using a homemade ELISA, with a hybridome provided by Dr Michel Tremblay as described previously^62^. Levels of IL-17A, IL-10, IL-22 and IFN-γ in cell culture supernatants were measured using commercial kits, according to the manufacturer’s instructions.

### Quantification of HIV-DNA

Early reverse transcripts (RU5) were quantified using SYBR Green PCR, while late reverse transcripts and integrated HIV-DNA levels were quantified by nested real-time PCR, as we previously reported^17^, using specific primers and probes indicated in Supplemental Table 4. Briefly, HIV-infected cells were digested at a concentration of 50,000 cells/15 µl of lysis buffer containing 0.1 mM Tris HCl pH 8.0 and 10 mg/mL proteinase K. HIV-DNA levels were measured relative to CD3-DNA used as an internal control for cell numbers (2 CD3-DNA copies per 1 cell). Serial dilutions from 3×10^5^ to 3 ACH-2 cells were used as a standard curve for absolute quantification. The limit of detection of HIV/CD3 was 3 copies per test.

### SYBR Green quantitative real time RT-PCR

One step RT-PCR was performed using SYBR Green dye, with the light Cycler 480 II (Roche, Basel, Switzerland). Total RNA was isolated with the Qiagen RNeasy Plus Mini Kit. A minimum of 10 ng RNA was loaded in each well in triplicate. The relative gene expression was normalized to the housekeeping gene 28S rRNA and quantitative gene expression was assessed using a cDNA standard obtained from RNA extracted from CD4^+^ T cells.

### Viral outgrowth assay (VOA)

A simplified VOA was performed, as we previously described^17,61,62^. Briefly, memory CD4^+^ T-cells were isolated by negative selection using magnetic beads from PBMCs of ART-treated PWH, activated with immobilized anti-CD3 and soluble anti-CD28 antibodies and cultured in 48-well plates (1×10^6^ cells/ml RPMI media, four replicates per condition) in presence of SR9011 or SR8278 (concentrations indicated in Figure legends). At day 3 post-TCR activation, cells were washed and split into two new 48-well plate wells and cultured in RPMI1640 media supplemented with rhIL-2 and SR9011/SR8278. Cells were further split at day 6 and 9 post-stimulation and half of the media was replenished with rhIL-2 containing SR9011/SR8278. Cells were harvested at day 12 and intracellular HIV-p24 expression was quantified by FACS. Soluble HIV-p24 levels were quantified by ELISA in supernatants harvested at days 3, 6, 9 and 12 post-TCR triggering.

### Western Blotting

To prepare samples for Western blot, 2×10^6^ TCR-activated CD4+ T cells per condition were cultured in 48-well plates for 3 or 6 days, washed with 1X PBS, pelleted down and frozen at - 80°C. Cells were lysed with RIPA buffer supplemented with phosphatase inhibitor and protease inhibitor cocktail for 5 minutes at 4 °C, sonicated and centrifuged at 13,000*g* for 10 minutes. Proteins were quantified with DC Protein Assay. 8 µg protein/well or condition were loaded on SDS PAGE gel electrophoresis and run for 75 minutes at 130 volts, then transferred on immobilon-PSQ polyvinylidene difluoride (PVDF) membrane for 70 minutes at 100 volts and blocked with 5% BSA. PVDF membrane was blocked with Tris-Buffered Saline (TBS) 0.1% Tween 5% Bovine Serum Albumin for 45 minutes and respectively incubated overnight with anti-SAMHD1 Thr592, total anti-SAMHD1, anti-APOBEC3G, anti-β-Tubulin and anti-β-actin antibodies. Immunoreactive bands were detected using HRP-conjugated secondary antibodies and revealed with ECL substrate on the Chemidoc imaging system from Bio-Rad.

### Illumina RNA Sequencing and Analysis

Cells were harvested and total RNA was extracted using the Qiagen All Prep DNA/RNA/miRNA Universal Kit. The genome-wide bulk RNA sequencing was performed by Génome Québec using the polyA enriched RNA (stranded) library and the Illumina Technology (NovaSeq PE100-25M Reads). The paired-end sequencing reads were aligned to coding and non-coding transcripts from Homo Sapiens database GRCh 37 version75 and quantified with the Kallisto software version 0.44.0. Statistical analyses were performed using R version 4.21. Differentially expressed genes (DEG) were identified based on p-values (p<0.05), adjusted p-values (adj. p<0.05) and fold-change (FC, cutoff 1.3). Differential expression analysis was performed using the limma Bioconductor R package (version 3.52.2) on the log2-counts per million (logCPM) transformed transcript-level and gene-level data. Gene set enrichment analysis (GSEA; C2, C3, C5, C7, C8, and Hallmark databases; https://www.gsea-msigdb.org/gsea/msigdb/index.jsp) was performed using the gene set variation analysis (GSVA) method (package version 1.344.2) on the logCPM data using a Gaussian cumulative distribution function.

To obtain the list of targets from transcription factors, a motif search was performed using PWM tools on the Eukaryotic Promoter Database^108^, using Position Weight Matrix (PWM) models (https://epd.expasy.org/pwmtools). Homo sapiens promoters were used as a reference for the motif search, with a search window ranging from −1kb to +100b from the transcription start site (TSS). The motifs used are BMAL1_HUMAN.H11MO.0.A and RORG_HUMAN.H11MO.0.C for BMAL/CLOCK and REV-ERB alpha/beta respectively, as found in the motif database HOCOMOCO v11^109^. The links to motifs are:

BMAL1 motif:

https://epd.expasy.org/cgi-bin/pwmtools/pwmviewer.cgi?ID=BMAL1_HUMAN.H11MO.0.A&lib=hocomocov11_human

RORC motif:

https://epd.expasy.org/cgi-bin/pwmtools/pwmviewer.cgi?ID=RORG_HUMAN.H11MO.0.C&lib=hocomocov11_human

The HIV targets list is a combination of two gene sets from the NCBI HIV-1 database based on the keywords “upregulates” and “interacts with”.

## STATISTICAL ANALYSES

### Statistical analysis

Statistical analyses were performed using the Prism software version 10.1.2 (GraphPad, Inc., La Joya, CA, USA). Statistical tests applied were Friedman (uncorrected Dunn’s test), Wilcoxon or Mann-Whitney test. P-values are indicated on the graphs with statistical significance for p<0.05.

## SUPPLEMENTAL TABLES

**Supplemental Table 1:** Clinical parameters of ART-treated PWH.

**Supplemental Table 2:** Clinical parameters of PWoH.

**Supplemental Table 3:** Key Resources.

**Supplemental Table 4:** Primers and probes used for PCR and RT-PCR.

## SUPPLEMENTAL FILES

**Supplemental Files 1**: Top differentially expressed genes in memory CD4+ T-cells of ART-treated PWH stimulated *via* the TCR in the presence of SR9011.

**Supplemental Files 2**: Top differentially expressed genes in memory CD4+ T-cells of ART-treated PWH stimulated *via* the TCR in the presence of SR8278.

**Supplemental File 3:** Transcripts modulated by SR8278 in memory CD4+ T-cells of ART-treated PWH included in the HIV interactom database and exhibiting BMAL binding motifs in their promoters

**Supplemental File 4:** Transcripts modulated by SR8278 in memory CD4+ T-cells of ART-treated PWH included in the HIV interactom database and exhibiting RORC binding motifs in their promoters

