## Supplemental Table 1 for "REV-ERBα/β Targeting Transcriptionally Reprograms HIV-Infected CD4^+^ T-Cells for Increased Viral Reactivation but Limited Virion Spread"

**Supplemental Table 1:** Clinical parameters of ART-treated PWH in the study.

| Participant ID | Sex | Age* | Ethnicity | CD4# | CD8# | CMV | PVL⊗ | Time since infection** | ART | Time of aviremia** |
|---|---|---|---|---|---|---|---|---|---|---|
| **ART #1** | M | 36 | C | 542 | 803 | - | <40 | 13 | Stribild | 12 |
| **ART #2** | M | 51 | C | 841 | 1,322 | + | <40 | 149 | Sustiva/ Truvada | 1 |
| **ART #3** | M | 30 | C | 598 | 605 | + | <40 | 80 | Stribild | 1 |
| **ART #4** | M | 47 | C | 425 | 1,156 | + | <40 | 182 | Atripla | 24 |
| **ART #5** | M | 64 | C | 649 | 620 | + | <40 | 186 | Triumeq | 1 |
| **ART #6** | M | 31 | L | 775 | 1,000 | + | <40 | 69 | Complera | 36 |
| **ART #7** | M | 54 | C | 886 | 579 | + | <40 | 60 | Truvada/ Isentress | 12 |

M, male; *, years; C, Caucasian; L, Latino-American; #, cells/µl; CMV, cytomegalovirus status; (-), negative; (+), positive; PVL, plasma viral load; ⊗, HIV-RNA copies per ml plasma; **, months; ART, antiretroviral therapy
