## Supplemental Table 2 for "REV-ERBα/β Targeting Transcriptionally Reprograms HIV-Infected CD4^+^ T-Cells for Increased Viral Reactivation but Limited Virion Spread"

**Supplemental Table 2:** Clinical parameters of PWH in the study.

| Participant ID | Sex | Age* | Ethnicity | CD4[#] | CD8[#] | CMV status |
|---|---|---|---|---|---|---|
| **Donor #1** | M | 69 | C | 517 | 186 | + |
| **Donor #2** | M | 26 | C | 302 | 127 | - |
| **Donor #3** | M | 47 | C | 604 | 147 | - |
| **Donor #4** | M | 34 | H | 468 | 707 | + |
| **Donor #5** | M | 59 | C | 551 | 198 | - |
| **Donor #6** | F | 43 | C | 572 | 246 | + |
| **Donor #7** | M | 60 | C | 704 | 487 | + |
| **Donor #8** | M | 63 | C | 700 | 344 | + |
| **Donor #9** | M | 40 | C | 1144 | 826 | + |
| **Donor #10** | M | 44 | C | 845 | 333 | + |
| **Donor #11** | M | 68 | C | 865 | 589 | + |
| **Donor #12** | M | 32 | C | 727 | 464 | - |
| **Donor #13** | M | 31 | C | 537 | 492 | + |
| **Donor #14** | M | 39 | C | 1424 | 1505 | + |
| **Donor #15** | M | 45 | C | 683 | 234 | + |
| **Donor #16** | F | 61 | C | 670 | 460 | + |
| **Donor #17** | M | 41 | C | 668 | 239 | + |
| **Donor #18** | M | 68 | C | 790 | 313 | + |

M, male; F, female; *, years; C, Caucasian; H, Hispanic #, cells/mm3; CMV, cytomegalovirus status; (-), negative; (+), positive
