## Supplemental Table 3 for "REV-ERBα/β Targeting Transcriptionally Reprograms HIV-Infected CD4^+^ T-Cells for Increased Viral Reactivation but Limited Virion Spread"

**Supplemental Table 3:** Key resources.

| Reagent or resource | Source | Identifier |
|---|---|---|
| Antibodies | | |
| Purified NA/LE Mouse Anti-Human CD3 | BD Pharmingen | Cat#555329 |
| Purified NA/LE Mouse Anti-Human CD28 | BD Pharmingen | Cat#555725 |
| Monoclonal mouse anti-SAMHD1 antibody clone OTI3F5 (30 µl) | Cedarlane | Cat#TA502024S |
| Polyclonal anti-SAMHD1 (PHOSPHO THR592) (0,02 mg) antibody | Cedarlane | Cat#8005-0.02MG |
| Monoclonal rabbit anti-APOBEC3G antibody | Cell Signaling Technology | Cat#43584S |
| Monoclonal mouse anti-®-actin antibody | Sigma | Cat#A5441 |
| Monoclonal rabbit anti-®-tubulin antibody | Cell Signaling Technology | cat#2146S |
| Goat Anti-mouse IgG (H+L) Secondary Antibody, HRP | Thermo Fisher | Cat#32340 |
| Goat anti-rabbit IgG HRP-linked Antibody | Cell Signaling | Cat#7074 |
| Mouse anti-human CD3 Pacific Blue | BD | Cat#558117 |
| Mouse anti-human CD4 Alexa Fluor 700 | BD | Cat#557922 |
| Mouse anti-human CD8 FITC | Miltenyi | Cat#130-113-157 |
| Mouse anti human CD45RA APCeFluor780 | eBioscience | Cat#47-0458-42 |
| Mouse anti-human CCR5 PE | BD | Cat#555993 |
| Mouse anti-human Ki-67 BUV395 | BD | Cat#564071 |
| Mouse anti-human Ki-67 PE | Biolegend | Cat#350504 |
| Mouse anti-human BST2 BV421 | BD | Cat#566381 |
| Mouse anti-human CD36 BUV737 | BD | Cat#748647 |
| Mouse Anti-Human TNF BV750 | BD | Cat#566359 |
| HIV-1 core (p24) antigen-RD1, KC57 | Beckman Coulter | Cat#6604667 |
| HIV-1 core (p24) antigen-FITC, KC57 | Beckman Coulter | Cat#6604665 |
| Anti-env Alexa Fluor 647 | Introgen | Cat#A21445 |
| PGT126 | From Dr. Finzi (obtained from International AIDS Vaccine Initiative, (IAVI)) | N/A |
| PGT151 | From Dr. Finzi (obtained from International AIDS Vaccine Initiative, (IAVI)) | N/A |
| p24 Enzyme-linked Immunosorbent Assay (ELISA) | Homemade. Hybridome provided by Dr. Michel J. Tremblay (Bounou et al., J Virol., 2002) | N/A |
| Bacterial and virus strains | | |
| NL4.3BaL HIV plasmid | From Dr. Dana Gabuzda (Dana-Farber Cancer Institute, Boston, MA, USA) | N/A |

| | | |
|---|---|---|
| VSV-G Plasmid | National Institution of Health (NIH) | Cat#ARP-4693 |
| NL4.3BaL⊗env GFP | National Institution of Health (NIH) | Cat#ARP-12637 |
| **Biological samples** | | |
| Leukapheresis collected from HIV- participants | N/A | N/A |
| **Chemicals, peptides, and recombinant proteins** | | |
| Sodium Bicarbonate ($NaHCO_3$) | Sigma | Cat#S5761 |
| Sodium Carbonate ($Na_2CO_3$) | Sigma | Cat#223530-500G |
| Thimerosal | Sigma | Cat#T5125-10G |
| Tween 20 | Fisher Scientific | Cat#BP337-500 |
| Triton X-100 | Sigma | Cat#X100-500mL |
| Trypan Blue | Thermo Fisher | Cat#15250061 |
| Bovine Serum Albumin (BSA) | Bioshop | Cat#ALB001.500 |
| Albumin Standard | Thermo Scientific | Cat#23209 |
| Streptavidin Horseradish Peroxidase (Strep-HRP) | Biosynth | Cat#65R-S104PHRP |
| TMB Peroxidase Substrate | Biosynth | Cat#42R-TB102 |
| Phosphoric Acid ($H_3PO_4$) | Sigma | Cat#PX0996 |
| Nonidet P-40 | Bioshop | Cat#NON505 |
| Tris HCl | Bioshop | Cat#TRS002.500 |
| Proteinase K | Invitrogen | Cat#25530015 |
| Molecular Grade Water | Wisent | Cat#809-115-CL |
| Trypsin-EDTA (0.25%), phenol red | Gibco | Cat#25200072 |
| X-tremeGENE HP DNA Transfection Reagent | Roche | cat#6366244001 |
| Methanol | Fisher Scientific | Cat#A454-4 |
| ReBlot Plus Strong Antibody Stripping Solution | Sigma | Cat#2504 |
| 4X Laemmli Sample Buffer | Bio-Rad | Cat#1610747 |
| 2-Mercaptoethanol | Sigma | Cat#63689-100ML-F |
| Precision Plus Protein Dual Color Standards | Bio-Rad | Cat#1610374 |
| Glycine | Bioshop | Cat#GLN001.10 |
| Sodium Chloride | Bioshop | Cat#SOD002.5 |
| Acrylamide/Bis-acrylamide 30% Solution | Bioshop | Cat#ACR010.500 |
| UltraPure Tris | Invitrogen | Cat#15504-020 |
| Sodium Dodecyl Sulfate (SDS) | Bioshop | Cat#SDS001.1 |
| N,N,N’,N’-Tetramethyl Ethylenediamine (TEMED) | Sigma | Cat#T8133 |
| Ammonium Persulfate (APS) | Bioshop | Cat#AMP001.25 |
| Radioimmunoprecipitation Assay Buffer (RIPA) 10X | Cell Signaling | Cat#9806S |
| Phosphatase Inhibitor (PhosSTOP) | Roche | Cat#4906845001 |
| Complete, Mini, EDTA-free protease inhibitor | Roche | Cat#11836170001 |
| Formaldehyde solution | Sigma | Cat#F1635 |
| Protein Assay Reagent A | Bio-Rad | Cat#500-0113 |
| Protein Assay Reagent B | Bio-Rad | Cat#500-0114 |
| Protein Assay Reagent S | Bio-Rad | Cat#500-0115 |
| Agarose, Biotechnology Grade | Bioshop | Cat#AGA002.500 |
| GelRed Nucleic Acid Gel Stain | Biotium | Cat#41003 |
| 50 bp DNA Ladder | Invitrogen | Cat#10416014 |

| | | |
|---|---|---|
| 100 bp DNA Ladder | Invitrogen | Cat#15628019 |
| Sodium Azide | Bioshop | Cat#SAZ001 |
| Fetal Bovine Serum (FBS) | Wisent | Cat#091-150 |
| Newborn Calf Serum (NBCS) | Wisent | Cat#075-350 |
| Dimethyl Sulfoxide (DMSO) | Sigma | Cat#34869-500 |
| RPMI 1640 Medium (RPMI) | Gibco | Cat#11875-093 |
| EDTA | Bioshop | Cat#EDT001.1 |
| Phosphate Buffered Saline (PBS) 1X | Gibco | Cat#20012-027 |
| Phosphate Buffered Saline (PBS) 10X | Wisent | Cat#311-12-LL |
| Penicillin-streptomycin | Gibco | Cat#15140122 |
| Lymphocyte Separation Medium (LSM) | Wisent | Cat#305-010 CL |
| Ethyl alcohol anhydrous | Commercial Alcohols | Cat#P016EAAN |
| Isopropyl alcohol | Bioshop | Cat#ISO920.4 |
| Recombinant Human IL-2 Protein | R&D Systems | Cat#202-IL-050 |
| SR9011 | Cayman Chemical | Cat#11930 |
| SR8278 | Sigma-Aldrich | Cat#S9576 |
| Opti-MEM | Life Technologies | Cat#31985070 |
| Commercial assays | | |
| RNeasy Plus mini kit (250) | Qiagen | Cat#74136 |
| AllPrep DNA/RNA/miRNA Universal Kit (50) | Qiagen | Cat#80224 |
| QuantiTect SYBR Green RT-PCR Kit | Qiagen | Cat#204243 |
| Taq DNA Polymerase | Invitrogen | Cat#18038067 |
| Wizard Plus Minipreps DNA Purification System | Promega | Cat#A7100 |
| EndoFree Plasmid Maxi Kit | Qiagen | Cat#12362 |
| EasySep Human Memory CD4+ T Cell Enrichment Kit | STEMCELL | Cat#19157 |
| All Prep DNA/RNA/miRNA Universal Kit | Qiagen | Cat#80224 |
| Clarity Max ECL Western Blotting Substrate | Bio-Rad | Cat#1705062S |
| Detergent Compatible (DC) Protein Assay | Bio-Rad | Cat#5000111 |
| Fixation/Permeabilization Kit | BD | Cat#554714 |
| LIVE/DEAD Fixable Aqua Dead Cell Stain Kit (405 nm excitation) | Invitrogen | Cat#L34957 |
| eBioscience Foxp3 / Transcription Factor Staining Buffer Set | Invitrogen | Cat#00-5523-00 |
| BD Cytofix/Cytoperm Fixation/Permeabilization kit | BD | Cat#554714 |
| Human IL-17A Uncoated ELISA kit | Invitrogen | Cat#88-7176-88 |
| Human IFN-© Uncoated ELISA kit | Invitrogen | Cat#88-7316-88 |
| Human IL-10 | R&D | Cat#DY217B-05 |
| Human IL-22 | R&D | Cat#DY782-05 |
| Human TFN-〈 | R&D | Cat#DY210-05 |
| Human CCL20/MIP3〈 | R&D | Cat#DY360-05 |
| Oligonucleotides | | |
| See Supplemental Table 1: Oligonucleotides | See Supplemental Table 1 : Oligonucleotides | See Supplemental Table 1 : Oligonucleotides |
| Software and algorithms | | |

| | | |
|---|---|---|
| FlowJo version 10.8.1 | BD | https://www.flowjo.com/ |
| GraphPad Prism version 10.1.2 | GraphPad | https://www.graphpad.com/ |
| Image Lab version 6.1.0 | Bio-Rad | https://www.bio-rad.com/en-ca/product/image-lab-software?ID=KRE6P5E8Z |
| Others | | |
| Pre-Separation Filters 30 µm | Miltenyi | Cat# 130-041-407 |
| Immobilon-PSQ Polyvinylidene Difluoride (PVDF) | Sigma | Cat#ISEQ00010 |
| Deoxynucleoside Triphosphates (dNTP) | Thermo Fisher | Cat#10297018 |
| LC480 probe master mix | Roche | Cat#4707494001 |
| 100 mM dNTP set | Invitrogen | Cat#10297-018 |
| 293T | ATCC | Cat#CRL-3216 |
| ACH-2 | NIH HIV Reagent Program | Cat#ARP-349 |

*N/A, Not applicable*
