## Supplemental Table 4 for "REV-ERBα/β Targeting Transcriptionally Reprograms HIV-Infected CD4^+^ T-Cells for Increased Viral Reactivation but Limited Virion Spread"

**Supplemental Table 4:** Primers and probes used for PCR and RT-PCR.

| Identification | Sequence | Provider | Identifier |
|---|---|---|---|
| **Primer AA55** | 5'-CGT CTA GAG ATT TTC CAC AC-3' | IDT | N/A |
| **Primer AA55M** | 5'-GCT AGA GAT TTT CCA CAC TGA CTA A-3' | IDT | N/A |
| **Primer M667** | 5'-CTA ACT AGG GAA CCC ACT G-3' | IDT | N/A |
| **Primer LM667** | 5'- ATG CCA CGT AAG CGA AAC TCT GGC TAA CTA GGG AAC CCA CTG-3' | IDT | N/A |
| **CD3 internal primer 1** | 5'- CCT CTC TTC AGC CAT TTA AGT A-3' | IDT | N/A |
| **CD3 internal primer 2** | 5'- GGC TAT CAT TCT TCT TCA AGG T-3' | IDT | N/A |
| **CD3 external primer 1** | 5'-ACT GAC ATG GAA CAG GGG AAG-3' | IDT | N/A |
| **CD3 external primer 2** | 5'- CCA GCT CTG AAG TAG GGA ACA TAT-3' | IDT | N/A |
| **Primer GagR** | 5'- AGC TCC CTG CTT GCC CAT A-3' | IDT | N/A |
| **Primer Alu1** | 5'- TCC CAG CTA CTG GGG AGG CTG AGG-3' | IDT | N/A |
| **Primer Alu2** | 5'- GCC TCC CAA AGT GCT GGG ATT ACA G-3' | IDT | N/A |
| **Primer LambdaT** | 5'- ATG CCA CGT AAG CGA AAC T-3' | IDT | N/A |
| **Primer SK29** | 5'- ACT AGG GAA CCC ACT GCT-3' | IDT | N/A |
| **Primer SK30** | 5'- GGT CTG AGG GAT CTC TAG-3' | IDT | N/A |
| **Probe LTR-LC** | 5'-LC640-CACTCAAGGCAAGCTTTATTGAGGC-3'-phosphate | TIB MolBiol | N/A |
| **Probe LTR-FL** | 5'-CACAACAGACGGGCACACACTACTTGA-3'-Flurescein | TIB MolBiol | N/A |
| **Probe CD3-P1** | 5'-GGCTGAAGGTTAGGGATACCAATATTCCTGTCTC-3'-Flurescein | TIB MolBiol | N/A |
| **Probe CD3-P2** | 5'-LC640-CTAGTGATGGGCTCTTCCCTTGAGCCCTTC-3'-phosphate | TIB MolBiol | N/A |
| **Primer 28S Forward** | 5′-CGA GAT TCC TGT CCC CAC TA-3′ | IDT | N/A |
| **Primer 28S Reverse** | 5′-GGG GCC ACC TCC TTA TTC TA-3′ | IDT | N/A |
| **RORC2 Forward** | 5'-CTG CTG AGA AGG ACA GGG AG-3' | IDT | N/A |
| **RORC2 Reverse** | 5'-AGT TCT GCT GAC GGG TGC-3' | IDT | N/A |
| **CCR5 Forward** | 5'-TCA AGT GTC AAG TCC AAT CTA TGA C-3' | IDT | N/A |
| **CCR5 Reverse** | 5'-TGC TCT TCA GCC TTT TGC AG-3' | IDT | N/A |
| **NR1D1** | NA | Qiagen | GeneGlobal ID: QT00000413 |
| **NR1D2** | NA | Qiagen | GeneGlobal ID: QT00008897 |
| **ARNTL** | NA | Qiagen | GeneGlobal ID: QT00011844 |
| **CXCR4** | NA | Qiagen | GeneGlobal ID: QT00223188 |
| **IL17A** | NA | Qiagen | GeneGlobal ID: QT00009233 |

*NA, information not available; N/A, Not applicable*
