## Supplemental Figures 1-10 for "REV-ERBα/β Targeting Transcriptionally Reprograms HIV-Infected CD4^+^ T-Cells for Increased Viral Reactivation but Limited Virion Spread"

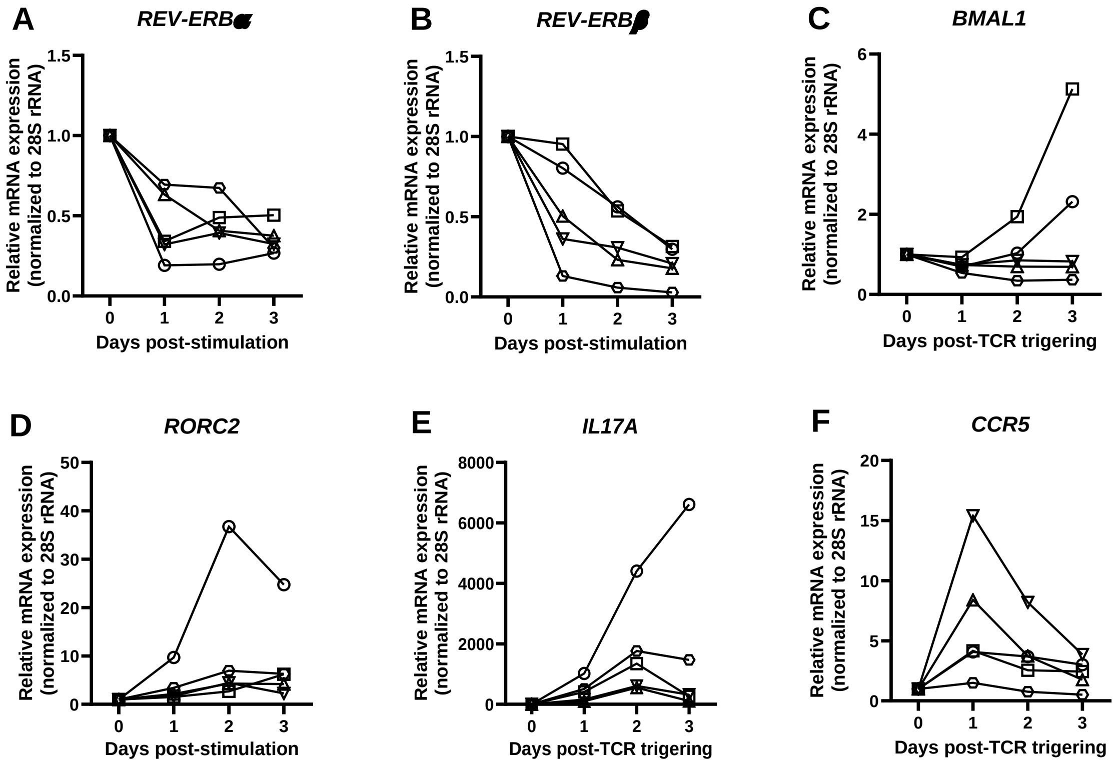


**Supplemental Figure 1: Kinetics of *REV-ERBα*, *REV-ERBβ*, *BMAL1*, *RORC2*, *IL17A*, and *CCR5* gene expression upon TCR triggering.** Memory $CD4^{+}$ T-cells were isolated from PBMCs of PWoH and activated *via* CD3/CD28 for up to 3 days. Cells at day 0 (*ex vivo*, no TCR triggering) or those collected at days 1, 2 and 3 post-TCR activation were used for total RNA extraction and real-time RT-PCR quantification of *REV-ERBα* **(A)**, *REV-ERBβ* **(B)**, *BMAL1* **(C)**, *RORC2* **(D)**, *IL17A* **(E)**, and *CCR5* **(F)** mRNA. Shown are the relative expressions of genes quantified and normalized to 28S rRNA levels, as a housekeeping gene. Depicted results are relative to day 0 (considered to be 1).

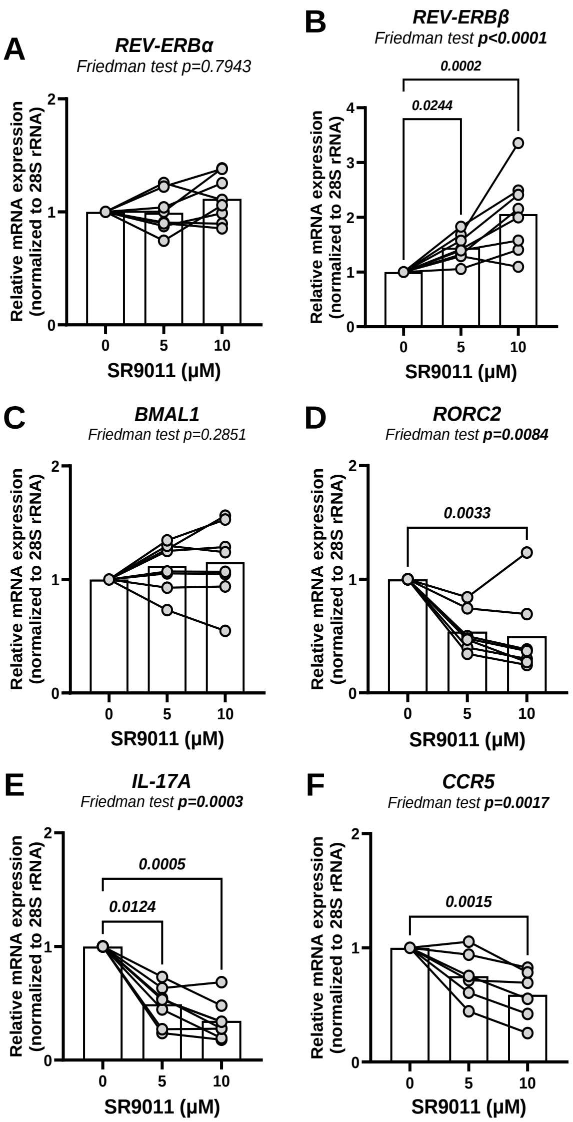


**Supplemental Figure 2: Effects of SR9011 on clock genes and HIV-1 co-receptor CCR5 mRNA expression.** Gene expression was quantified in memory CD4+ T-cells were isolated from PBMCs of PWoH, as described in Figure 2 legend, upon TCR triggering in the presence or the absence of SR9011 (5 and 10 μM), with DMSO used as control. **(A-F)** The levels of *REV-ERBα* and *REV-ERBβ* mRNA **(A-B)**, their target genes such as *BMAL1*, *RORC2* and *IL17A* (***C-E***), as well as *CCR5* mRNA **(F)**. RT-PCR quantification was performed at the time points described in Figure 2 legend. Gene expression was measured in triplicate by one-step real-time RT-PCR relative to 28S rRNA, as a reference gene. Shown are the values of relative gene expression relative to the DMSO condition. Friedman p-values and uncorrected Dunn's post-test p-values are indicated on the graphs. Statistically significant p-values (<0.05) are indicated in bold.

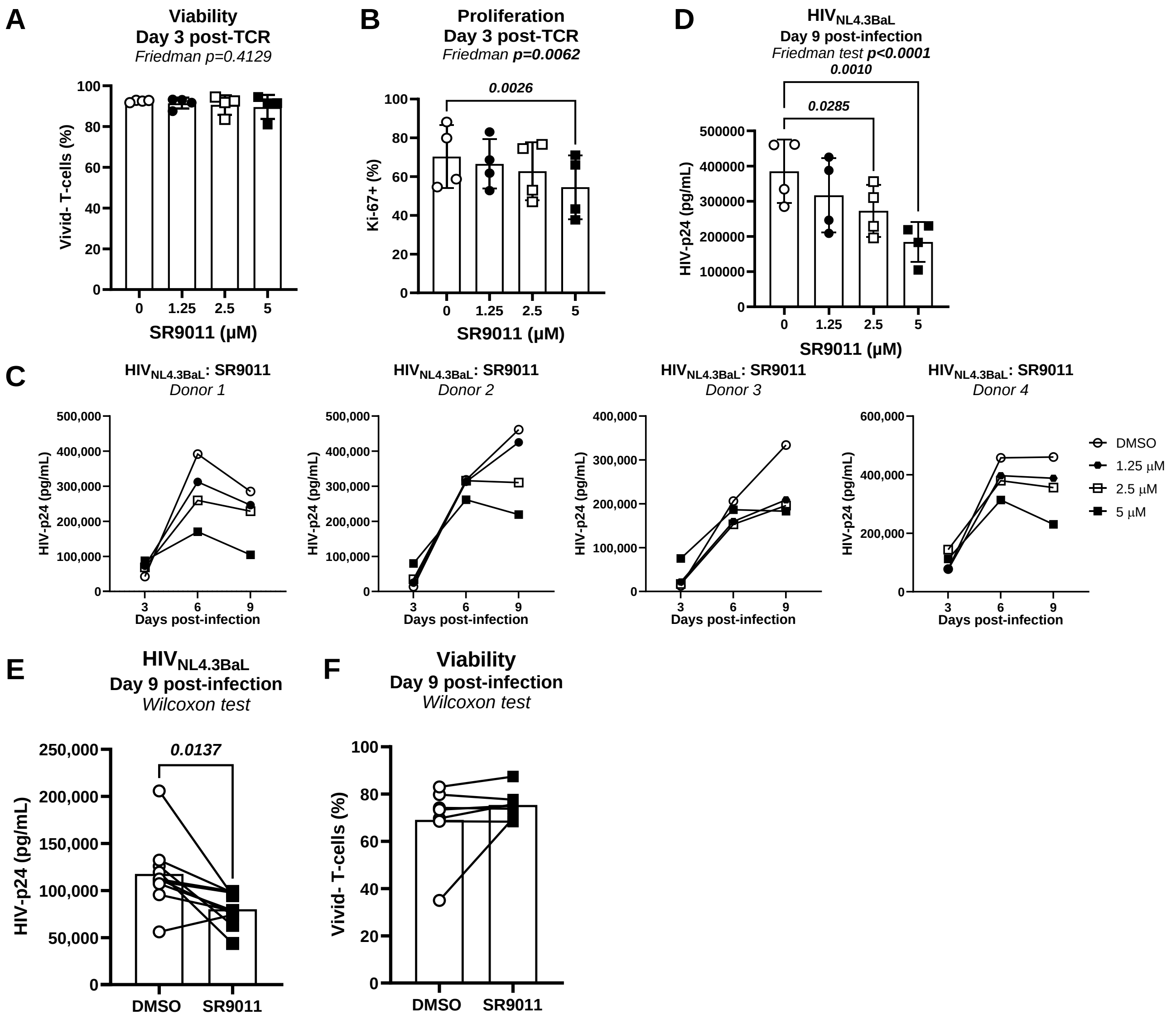


**Supplemental Figure 3: Dose-response studies on the effect of SR9011 on cell viability, proliferation and HIV-1 replication *in vitro*.** Memory CD4⁺ T-cells were isolated from PBMCs of PWoH, activated *via* CD3/CD28 for 3 days, exposed to $HIV_{NL4.3BaL}$ (25 ng/$10^6$ cells), and cultured with rhIL-2 (5 ng/ml) in the presence/absence of SR9011 (1.25; 2.5; 5 μM) for up to 9 additional days. **(A-D)** Show the statistical analysis of cell viability **(A)** and intra-nuclear Ki-67 expression **(B)** measured by flow cytometry at day 3 post-TCR activation, as well as the kinetics of HIV-1 replication in 4 individual participants at days 3, 6 and 9 post-infection **(C)**, and the statistical analysis of the effects of SR9011 on HIV-p24 levels measured by ELISA in cell culture supernatants harvested at day 9 post-infection (n=4) **(D)**. Friedman p-values and uncorrected Dunn's post-test p-values are indicated on the graphs. **(E-F)** Similar experiments were performed with $CD4^+$ T-cells from n=10 different HIV- participants infected with HIV-1 in the presence of SR9011 at 5 μM. Shown are statistical analysis of HIV-p24 levels measured by ELISA in cell culture supernatants **(E)**, and cell viability measured by flow cytometry at day 9 post-infection **(F)**. Wilcoxon test p-values are indicated on the graphs.

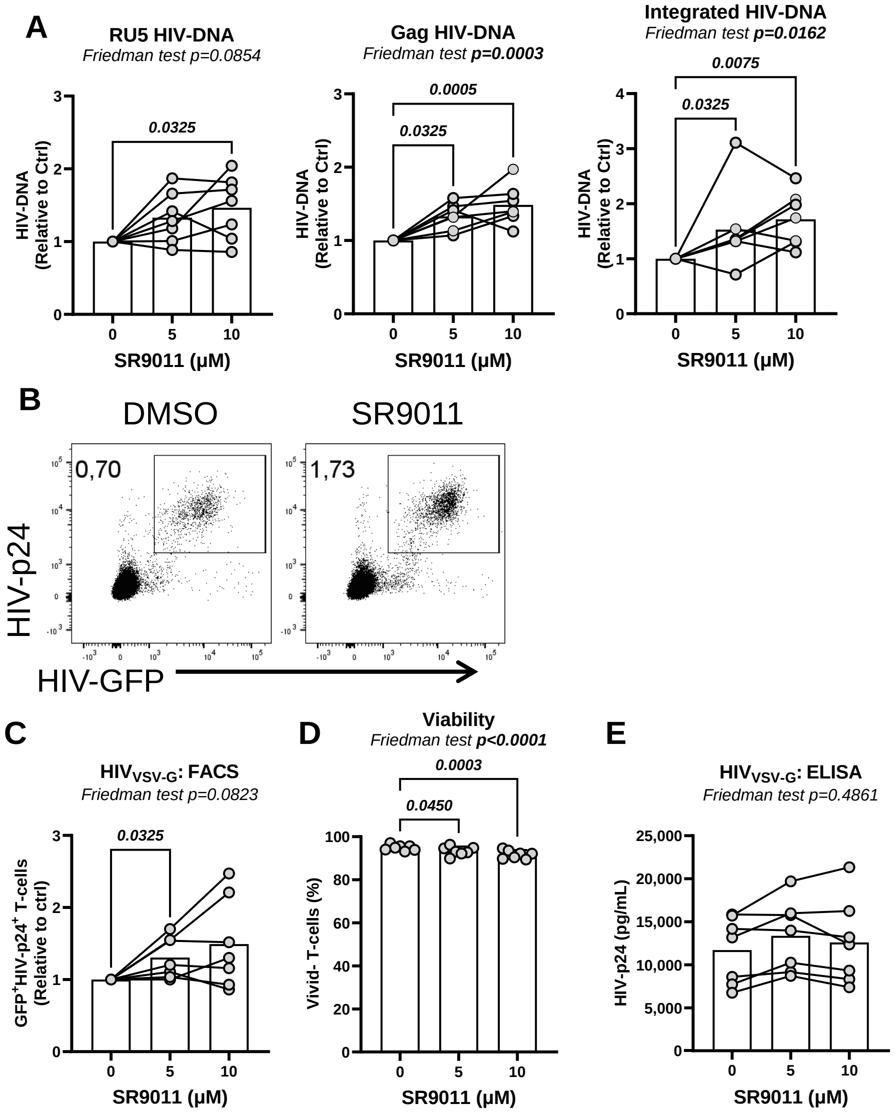


**Supplemental Figure 4: Effects of SR9011 on HIV-1 reverse transcription, integration, translation and virion release.** Experiments were performed as depicted in Figure 4A. Briefly, memory CD4+ T-cells were isolated from PBMCs of PWoH, stimulated with CD3 and CD28 Abs for 3 days, exposed to single-round VSV-G-pseudotyped HIV ($HIV_{VSV-G}$,100 ng/$10^6$ cells), and cultured with rhIL-2 (5 ng/ml) for 3 days. SR9011 (5 and 10 µM) was added to cell cultures immediately after infection. **(A)** Shown are levels of early (RU5), late (Gag) reverse transcripts and integrated HIV-DNA (copies/$10^6$ cells) measured by SYBR Green (RU5) and nested real-time PCR (Gag, Integrated) in CD4+ T-cells cultured in the presence/absence of SR9011 relative to DMSO. **(B-D)** Show the levels of intracellular HIV-p24 and GFP expression measured by flow cytometry for one representative donor **(B)**, as well as statistical analysis of intracellular HIV-p24 expression **(C)** and cell viability **(D)** (n=7). **(E)** Shows the levels of HIV-p24 in cell culture supernatants measured by ELISA at day 3 post-infection. Friedman and uncorrected Dunn's post-test p-values are indicated on the graphs. Statistically significant p-values (<0.05) are indicated in bold.

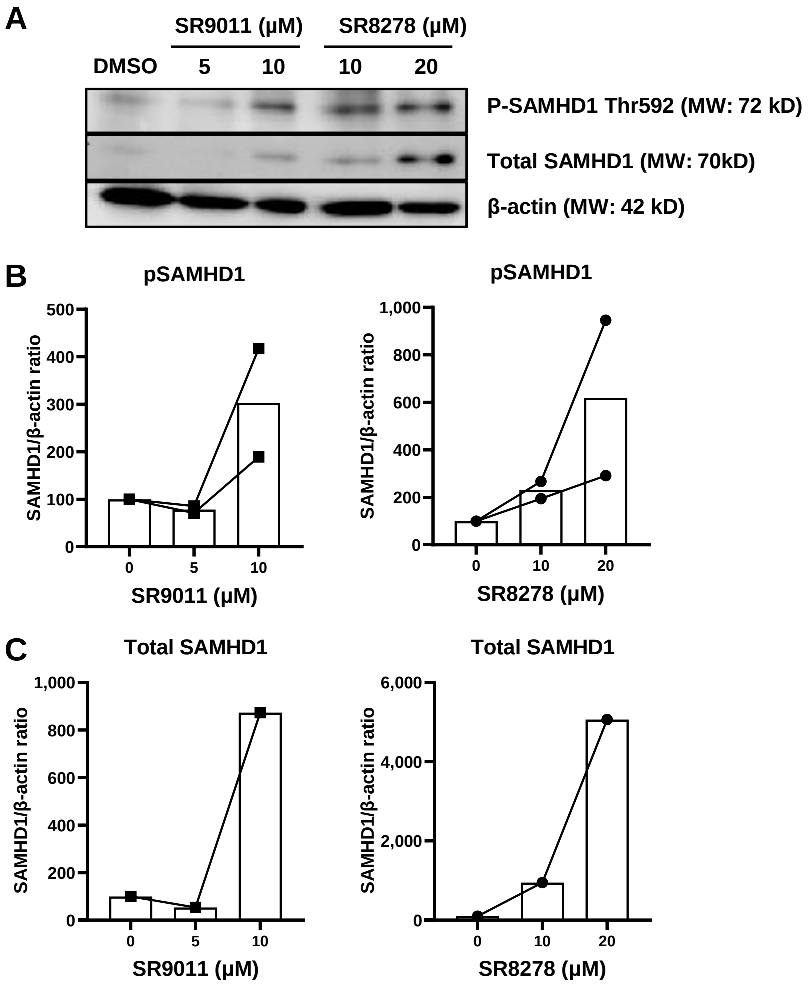


**Supplemental Figure 5: Effect of SR8978 and SR9011 on SAMHD1 expression and phosphorylation.** Memory CD4⁺ T-cells enriched from PBMCs of PWoH were activated via CD3/CD28 for 3 days (1x10$^6$ cells/well). Media was refreshed and cells were further cultured with rhIL-2 (5 ng/ml) in the presence/absence of SR9011 (5 and 10 µM) or SR8278 (10 and 20 µM) for an additional 24 hours. **(A)** Shows the western blot images from one representative participant for SAMHD1 phosphorylated (P) at Thr592 residue, total SAMHD1, and β-actin. **(B-C)** The density of the bands was measured. Shown is the ratio between the density of the P-SAMHD1 or total SAMHD1 and β-actin bands in cells exposed to SR9011 or SR8278 *versus* DMSO in n=2 donors. The molecular weight (MW) is indicated on the graphs.

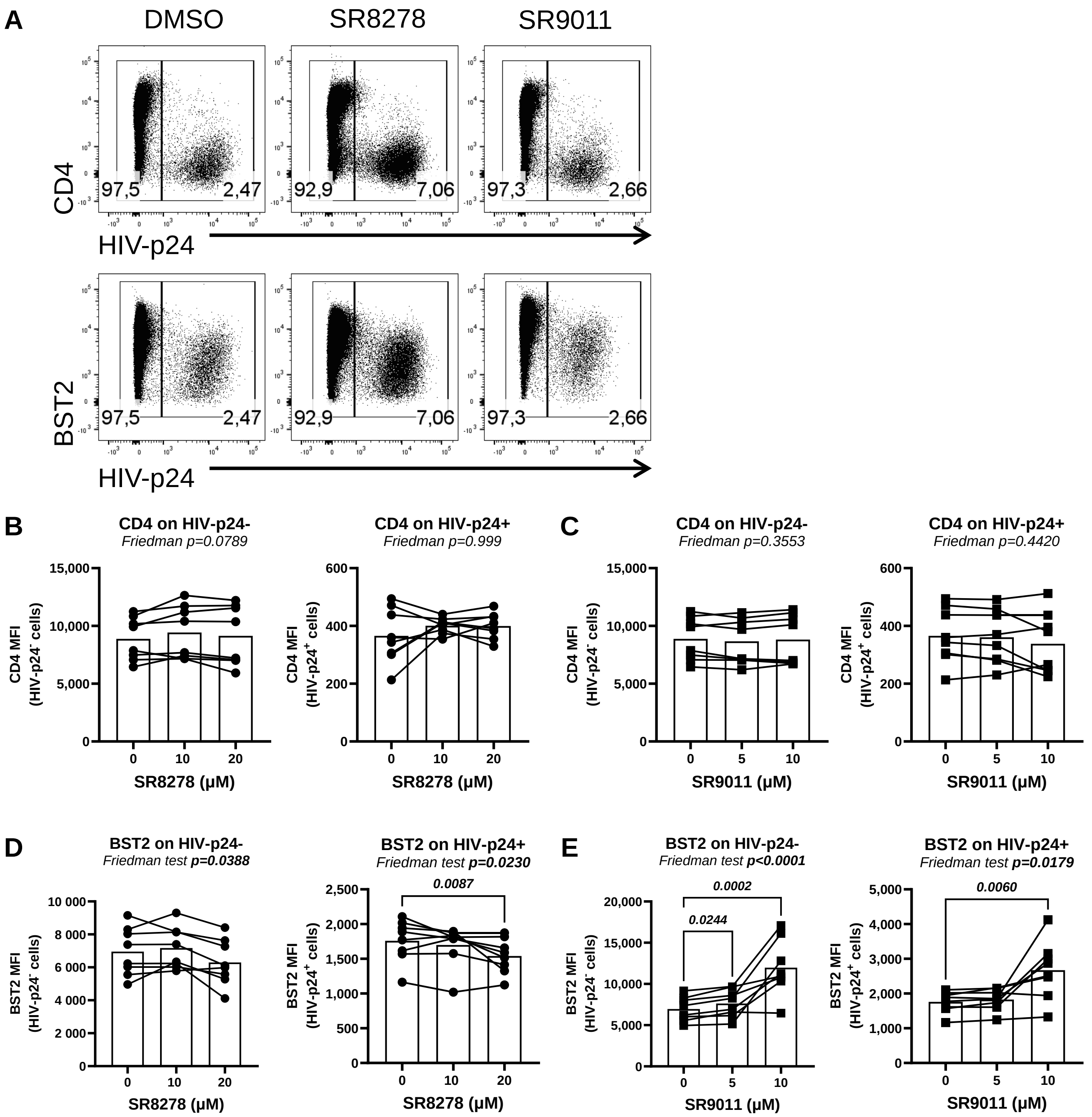


**Supplemental Figure 6: Effects of SR8978 and SR9011 on BST2 expression.** Memory $CD4^+$ T-cells isolated from PBMCs of PWoH were activated *via* CD3/CD28 for 3 days, exposed to single-round VSV-G-pseudotyped HIV (100 ng/$10^6$ cells), washed and cultured with rhIL-2 (5 ng/ml) in the presence/absence of SR8278 (10 and 20 µM) or SR9011 (5 and 10 µM) for 3 days. **(A)** Shows surface CD4 (upper panel) and BST2 (bottom panel) expression and intracellular HIV-p24 expression measured by flow cytometry for one representative donor. **(B-E)** Show the statistical analyses of median CD4 **(B-C)** or BST2 MFI **(D-E)** in HIV-negative (HIV-$p24^-$) and productively infected cells (HIV-$p24^+$) at day 3 post-infection. Friedman and uncorrected Dunn's test p-values are indicated on the graphs.

Ngassaki Yoka et al., Supplemental Figure 6
Related to Figures 4

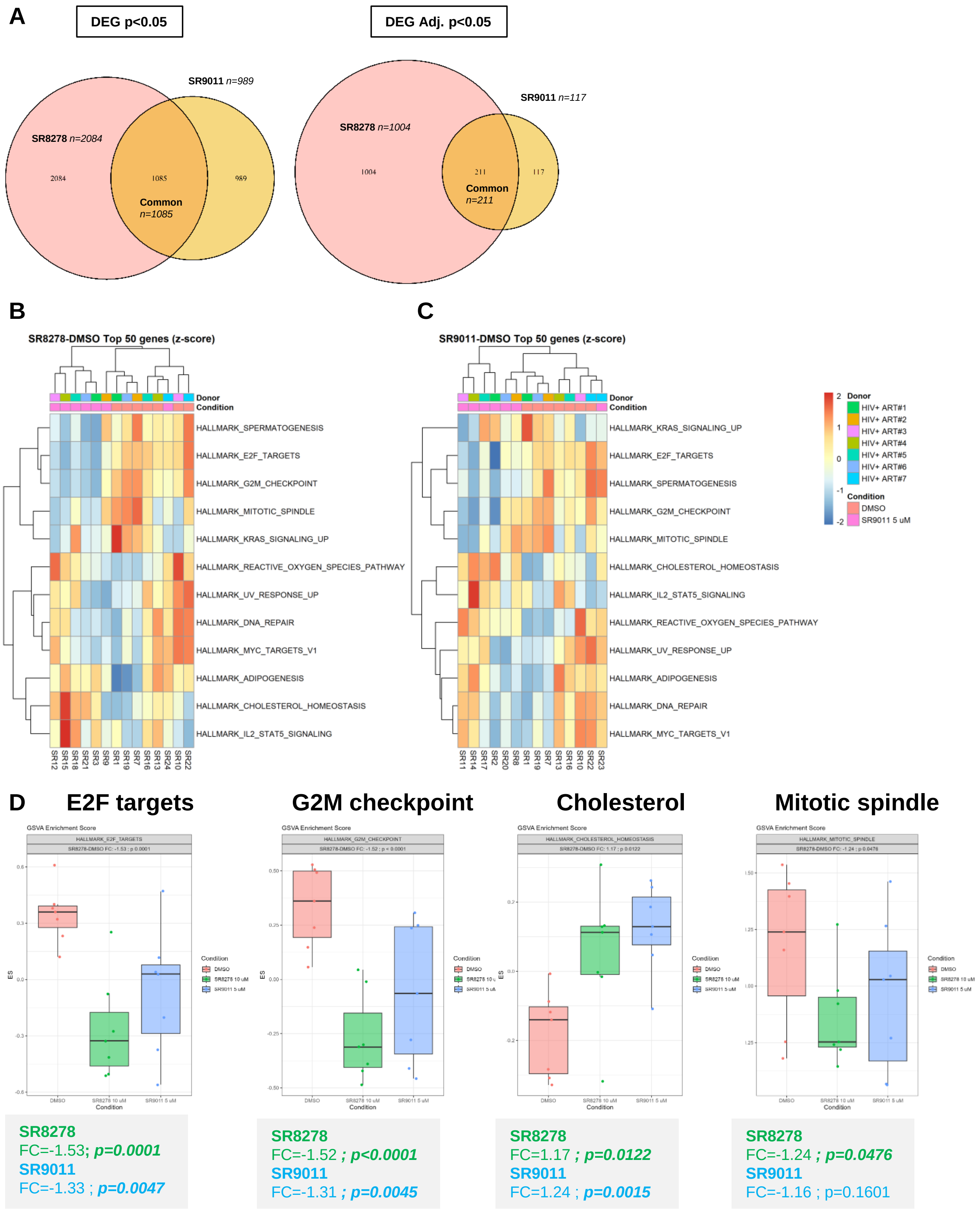


Ngassaki Yoka et al., Supplemental Figure 7
Related to Figure 6

**Supplemental Figure 7: GSVA reveals top modulated pathways by SR8278 and SR9011 in CD4+ T-cells of ART-treated PWH**. The genome-wide bulk RNA sequencing and GSVA were performed, as described in Figure 6 legend. **(A)** Shows the Venn diagrams depicting the number of DEGs modulated by SR8278 (10 µM) and SR9011 (5 µM), either unique or common. **(B-C)** Show the heatmaps depicting top Hallmark pathways modulated by SR8278 **(B)** and SR9011 **(C)**. **(D)** Shows the box plot graphs including the four top modulated pathways identified in GSVA based p-values < 0.05: E2F targets (left), G2M checkpoint (middle left), Mitotic spindle (middle right), and Cholesterol (right). Fold changes (FC) and p-values are indicated below the box plot graphs.

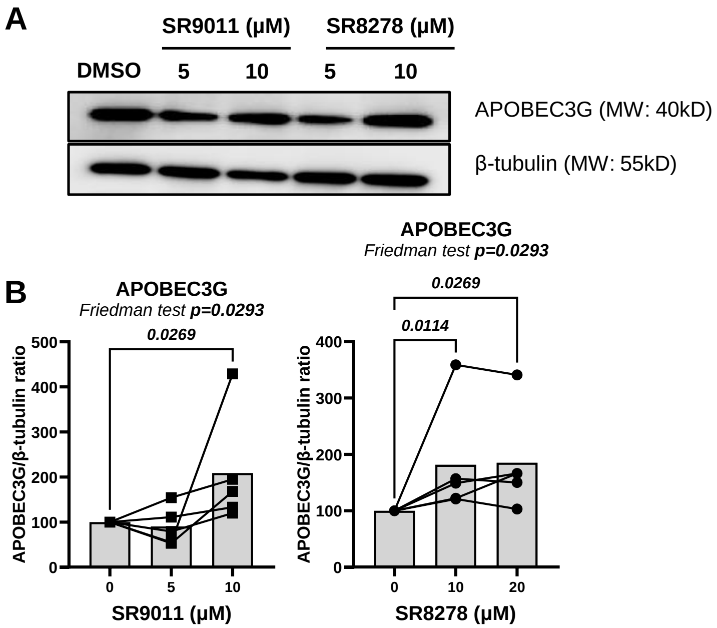


**Supplemental Figure 8: Effect of SR8978 and SR9011 on APOBEC3G expression.** Memory CD4+ T-cells enriched from PBMCs of ART-treated PWH were activated *via* the TCR and cultured in the presence/absence of SR9011 (5 and 10 μM) or SR8278 (10 and 20 μM) for one day. **(A)** Shows the Western blot results from one representative donor for APOBEC3G and β-tubulin. The density of the bands was measured. **(B)** Shows the ratio between APOBEC3G and β-tubulin band density in CD4+ T-cells cultured in the presence/absence of SR9011 or SR8278. Friedman p-values and uncorrected Dunn's post-test p-values are indicated on the graphs. The molecular weight is indicated on the graphs.

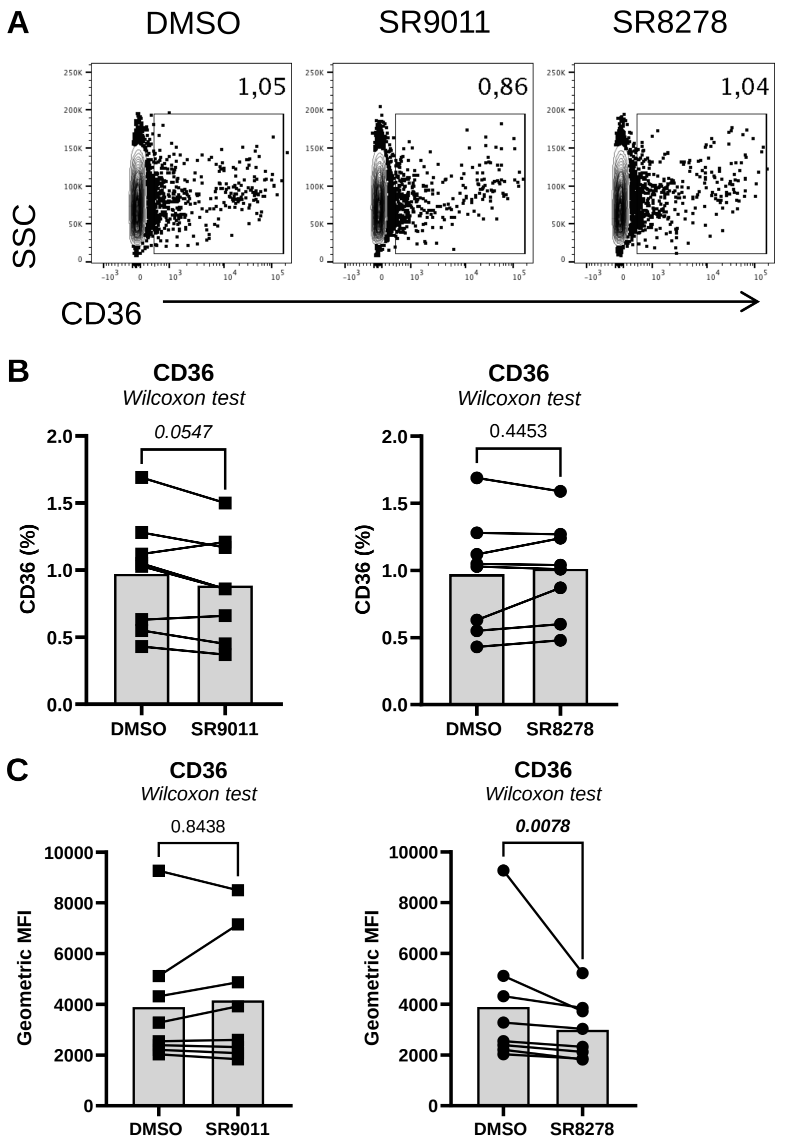


**Supplemental Figure 9: Effect of SR8978 and SR9011 on surface CD36 expression.** Memory CD4+ T-cells enriched from PBMCs of ART-treated PWH were activated *via* CD3/CD28 for 2 days (2x$10^6$ cells/well) and cultured in the presence/absence of SR9011 (5 µM) and SR8278 (10 µM). Then, cells were harvested and stained on the surface with CD36 Abs and analyzed by flow cytometry. **(A)** Shows CD36 surface expression for one representative donor. **(B-C)** Show the statistical analyses of the effect of SR9011 and SR8278 on the % **(B)** and MFI **(C)** expression of CD36 on CD4+ T-cells from n=8 ART-treated PWH. Wilcoxon test p-values are indicated on the graphs.

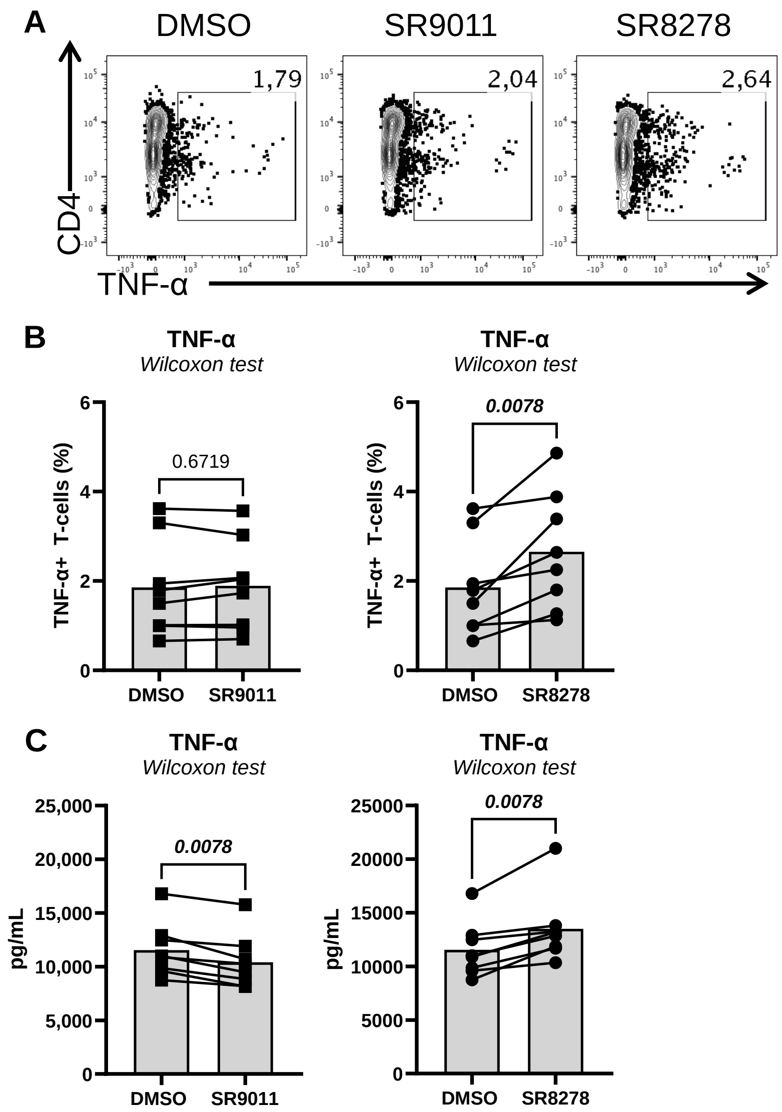


**Supplemental Figure 10: Effects of SR8978 and SR9011 on TNF-α intracellular expression and extracellular release.** Memory CD4+ T-cells enriched from PBMCs of ART-treated PWH were activated *via* CD3/CD28 for 2 days (2x$10^6$ cells/well) and cultured in the presence/absence of SR9011 (5 µM) and SR8278 (10 µM). **(A-B)** Cells were harvested and stained intracellularly with fluorochrome-conjugated TNF-α Abs and analyzed by flow cytometry. Shown are dot plots of intracellular TNF-α expression for one representative donor **(A)** and statistical analysis of the effect of SR9011 and SR8278 on intracellular TNF-α expression in CD4+ T-cells from n=8 ART-treated PWH **(B)**. **(C)** Shows TNF-α levels measured by ELISA in cell culture supernatants harvested from experiments performed with CD4+ T-cells from n=8 ART-treated PWH.
